# Periplasmic gatekeeping of phage DNA entry by rSAM enzyme-mediated maturation of an effector with His-X-Ser repeats

**DOI:** 10.64898/2026.01.06.697848

**Authors:** Mengling Li, Erchao Sun, Shuangshuang Wang, Yuxuan Huang, Haiguang Song, Sirong Kuang, Guoqi Li, Yuepeng Liu, Ziwei Xia, Xueqi Zhang, Jiumin Han, Zhaoxu Dong, Venigalla B. Rao, Lvhui Sun, Pan Tao

**Affiliations:** State Key Laboratory of Agricultural Microbiology, Cooperative Innovation Center for Sustainable Pig Production, College of Veterinary Medicine, Huazhong Agricultural University, Wuhan, Hubei 430070, China; Hubei Hongshan Lab, Wuhan, Hubei 430070, China; Bacteriophage Medical Research Center, Department of Biology, The Catholic University of America, Washington, DC 20064, USA

## Abstract

In the relentless arms race between bacteria and phages, bacteria have evolved a variety of defense strategies to combat phage infection. However, no bacterial system has been identified that specifically target the phage DNA entry phase. Here, we present a novel bacterial antiphage system, termed HXS, which provides broad-spectrum and robust antiphage activity by interfering with phage DNA entry. The HXS system consists of two radical S-adenosylmethionine (rSAM) enzymes (HxsB and HxsC), a small protein (HxsD), and the effector HxsA with a peptidoglycan-binding domain and five His-Xaa-Ser (HXS) repeats. HxsB/HxsC catalyze rSAM enzyme-dependent maturation of HxsA, including N-terminal processing and a site-specific +8 Da modification, thereby producing a periplasmic effector required for HXS defense. Biochemical evidence supports a model in which the matured effector likely engages incoming DNA electrostatically to arrest entry, establishing an rSAM enzyme-modified protein effector in antiphage defense.

## INTRODUCTION

Bacteria and phages are among the most abundant and dynamic biological entities on Earth, engaged in a continuous evolutionary arms race that profoundly shapes microbial diversity and ecosystem function ^1,2^. Phages initiate their infection by attaching to the bacterial surface receptors and injecting their genomic DNA into host cells. After DNA replication, transcription, and then translation, the phage progenies are assembled and finally released, usually resulting in the lysis of host cells ^3^. Recent studies have shown that bacteria have evolved a variety of antiphage systems to thwart different stages of the phage replication cycle to counteract phage predation and ensure survival ^4-6^.

As a first line of defense, bacteria can alter the phage receptors on their cell surfaces, such as surface proteins or lipopolysaccharides (LPS), by mutation or chemical modification to prevent phage recognition and binding ^7-9^. After phage attachment and DNA entry, bacteria use systems such as restriction-modification (RM) and CRISPR-Cas to identify and degrade phage DNA to protect themselves ^10,11^. During phage DNA replication, transcription, translation, and virion assembly, a variety of antiphage systems such as CBASS ^12,13^, Thoeris ^14,15^, TIR ^16^, and toxin-antitoxin ^17,18^ can be activated, usually resulting in premature host cell death or abortion of the infection process, thereby preventing assembly and release of mature phage progeny.

Although many well-characterized bacterial defenses act after phage genomes have entered the cytoplasm, surprisingly, no bacterial system has yet been identified that specifically targets the phage genomic DNA entry step. While the superinfection exclusion (Sie) systems of T4 phage can prevent subsequent infection by other T-even-like phages by blocking the entry of phage DNA, these systems are phage-encoded self-protection with a narrow spectrum, acting mainly against closely related phages ^19^.

Here, we present a broad and robust bacterial antiphage system, which we call the HXS system, composed of four proteins that work together to uniquely inhibit the phage DNA entry. HxsB and HxsC have a radical S-adenosylmethionine (rSAM) domain, which is part of a diverse family of enzymes known for their roles in various biochemical processes, particularly the modification of peptides or proteins ^20^. HxsA contains five His-Xaa-Ser (HXS) repeats, a motif that has been identified in bacterial proteins for over a decade but whose function has remained largely mysterious^21^. We showed that the HxsA is cleaved by the coordinated action of HxsB, HxsC, and HxsD into a functional shorter and covalently modified protein that was translocated into periplasm to inhibit phage DNA entry. The HXS system combines the rSAM domains and HXS repeats and presents a novel function of these elements in the context of antiphage activity. Our findings reveal a previously unrecognized bacterial defense strategy, expanding our understanding of microbial interactions and providing a potential new target for biotechnological applications in phage therapy and bacterial resistance management.

## RESULTS

### The HXS system is a novel defense system with a broad antiphage activity

The HXS system was serendipitously discovered while we were trying to identify homologs of viperin proteins in *Escherichia coli* by sequence and structural domain comparisons. Mammalian viperin is an interferon-induced antiviral protein belonging to the rSAM superfamily ^22^. It produces the ribonucleotide analog, 3’-deoxy-3’,4’-didehydro-CTP (ddhCTP), which acts as a chain terminator during viral RNA synthesis, thereby inhibiting viral replication ^22^. A recent study has shown that a prokaryotic viperin also has antiphage activity that protects against T7 phage infection by inhibiting viral polymerase-dependent transcription, like mammalian viperin ^23^.

During the identification of viperin homologs in the *E. coli* genomes, we identified 114 proteins belonging to the COG0535 rSAM superfamily. These proteins were clustered into 2 clusters, with 80-90% amino acid sequence identity within each cluster. Interestingly, we found that nearly half of these proteins are located within a unique operon structure. This operon contains twin-like rSAM-containing proteins (belong to two 2 clusters respectively) in proximity, accompanied by two additional proteins (Fig. 1a). We cloned one representative operon into the pSEC1 vector and transformed it into *E. coli* MG1655 cells to assess its antiphage activity against a taxonomically diverse panel of coliphages, which we taxonomically classified into multiple ICTV-recognized genera based on their genome sequences (Supplementary Table 1). HXS inhibited 110 of 113 phages (97.3%), with most exhibiting ≥10^3-fold reductions in efficiency of plating (EOP) (Fig. 1a). The only recurrent exceptions were three phages: two Tequatrovirus (including canonical T4) and one Punavirus (the classical generalized transducing phage P1). These results indicate broad and potent antiphage activity across diverse coliphage lineages.

**Fig. 1.**
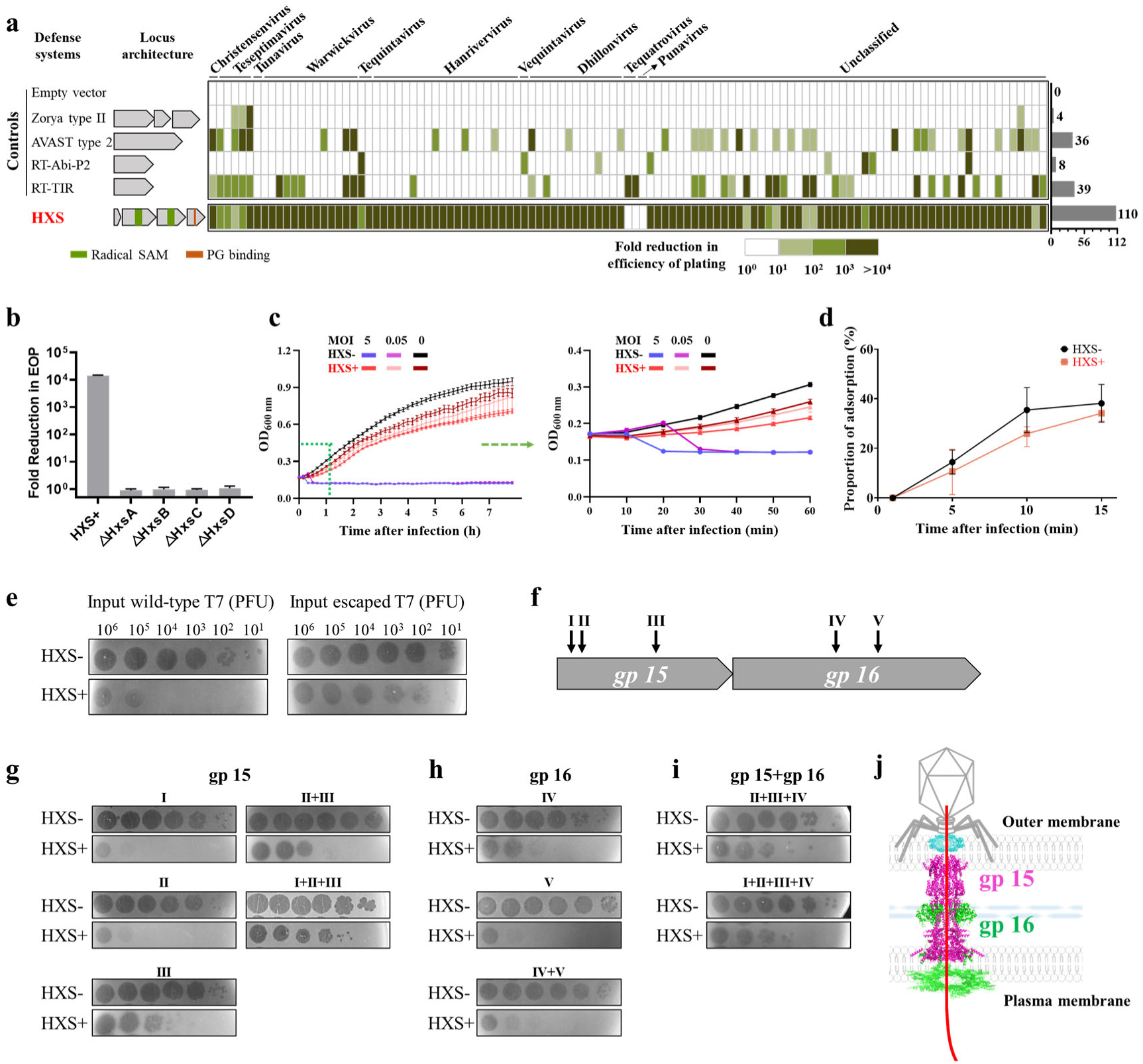
The HXS is a novel defense system with a broad antiphage activity. **a**, A heat map showing the antiphage activity of the HXS and 4 control defense systems against different phages. *E. coli* MG1655 cells expressing a defense system or the empty vector were infected individually with phages. The domain organization of the HXS systems was shown. The bar graph on the right side shows the number of phages restricted in each system. **b**, Single gene deletion analysis of the HXS. Wild-type and the mutated HXS were individually transformed into *E. coli* cells, which were infected with T7 phage. Empty vector was used as a control. **c**, Growth curves of *E. coli* cells expressing HXS after infection with T7 phage at different multiplicity of infection (MOI=0, 0.05, and 5). *E. coli* cells containing the empty vector were used as controls. **d**, Phage absorption assay in the presence or absence of HXS. Data were presented as the mean of three independent assays±SD. **e**, Plaque assays showing that the escaped T7 mutant has evolved the ability to escape HXS after 25 rounds of selection. Serial dilutions of each phage were spotted onto lawns of *E. coli* carrying either an empty vector or the HXS system. “Input phage (PFU)” denotes the number of PFU added per spot; 10^1^ to 10^6^ indicate the number of PFU added per spot in each condition. **f**, Schematic diagram showing the 5 mutation sites of the escaped T7 phage in the *gp15* and *gp16* genes, identified by sequence alignment with the wild-type genome (three mutations in *gp15* and two in *gp16*). Arrows indicate the location of the mutation sites. **g-i**, Identification of the critical mutation sites in the *gp15* and *gp16* genes. Single or multiple site mutations were introduced into the in the *gp15* and *gp16* genes of the wild-type T7 phage. The efficiency of these mutants to escape HXS was evaluated using plaque assays. **j**, A schematic diagram showing the T7 DNA-ejectosome composed of the gp15 and gp16 during infection. Gp15 (purple) assembles the hexameric translocation tunnel, and gp16 (green) serves as a “molecular tape” that stabilizes gp15-gp15 interfaces.

Single gene deletion experiments showed that all the four genes are necessary for the antiphage activity of the HXS system (Fig. 1b). By surveying 4,541 *E. coli* genomes available as of November 2025, we found that 77% of detected *hxsA-D* operons did not overlap with prophage regions predicted by geNomad v1.8.0 ^24^, indicating that HXS is typically chromosomally encoded rather than part of prophage cargo (Supplementary Fig. 1a-b). Bacterial growth curves showed that a high dose of phage infection (MOI=5) did not induce the rapid death of cells containing the HXS system (Fig. 1c), which is usually a feature of the abortive infection mechanism of many defense systems ^25-27^. The phage adsorption assay showed that the proportion of phages adsorbed onto *E. coli* in the presence of the HXS system did not change significantly at different time points after infection (Fig. 1d). These results indicated that the HXS system did not inhibit phage adsorption.

### The HXS system inhibits phage genomic DNA entry into *E. coli*

To obtain escape variants for probing how HXS blocks infection, we serially passaged phage T7 on HXS-expressing *E. coli*. T7 is a strictly lytic, genetically tractable, well-characterized coliphage that is robustly restricted by HXS. These features make it a practical and informative starting point for dissecting how HXS affects phage infection. After 25 passages, we recovered escape mutants capable of overcoming HXS (Fig. 1e; Supplementary Fig. 2). Genome sequencing of a single-plaque-purified escape clone identified seven mutations within the same phage genome (Supplementary Table 2), including five in the internal core proteins gp15 and gp16 (lower-case “gp” denoting “gene product”) (Fig. 1f), one synonymous mutation in gene *2*, and one intergenic mutation between gene *0.3* and gene *19.5*. We then introduced the single or multiple of these mutations into the wild-type T7 to generate ten T7 phage mutants as shown in Fig. 1g-i. Plaque assays showed that single mutations in the gp15 and the single or double mutation in the gp16 had a limited effect on T7 phage escape from the HXS system, whereas double or triple mutations in gp15 significantly enhanced the ability of T7 phage to escape from the HXS system. The gp15 and gp16 are key components of the phage T7 DNA-ejectosome that initiates genome translocation into the host cell (Fig. 1j) ^28-30^ ^31^. Gp15 assembles the hexameric translocation tunnel, and gp16 serves as a ‘molecular tape’ that stabilizes gp15-gp15 interfaces. Together, the mutation results suggest that the HXS may interfere with phage DNA entry.

T7 phage translocate its genomic DNA from the left end (highlighted in red in Fig. 2a and the fragment I in Fig. 2b) into *E. coli* cell ^32^. Based on the polarity of T7 phage DNA entry and the character of the endonuclease *Dpn*I that can specifically cleave fully methylated but not the unmethylated T7 genomic DNA, previous studies had developed the *Dpn*I digestion assay to directly measure the DNA translocation rate ^33^. We then used this assay to determine whether the HXS system interfered with T7 DNA translocation. Plaque assays showed that overexpression of Dam methylase did not affect the antiphage activity of HXS (Supplementary Fig. 3). The genomic DNA of T7 (containing 6 *Dpn*I sites) propagated in Dam methylase-overexpressing cells became methylated and was cleaved into 5 large fragments by *Dpn*I (Fig. 2b, right agarose gel). However, the T7 DNA prepared with standard *E. coli* is incompletely methylated and is *Dpn*I-resistant (Fig. 2b, left agarose gel). During infection of Dam-overexpressing cells by unmethylated T7, the left end that firstly enters the cytoplasm becomes methylated by Dam, whereas the capsid-retained portion remains unmethylated (Fig. 2a). The partially methylated phage DNA was isolated and digested by *Dpn*I. The appearance of *Dpn*I-digested DNA bands, determined by Southern blot with biotin-labeled probes specific for fragments I and V, over the infection course reflects the rate of T7 DNA translocation ^33^.

**Fig. 2.**
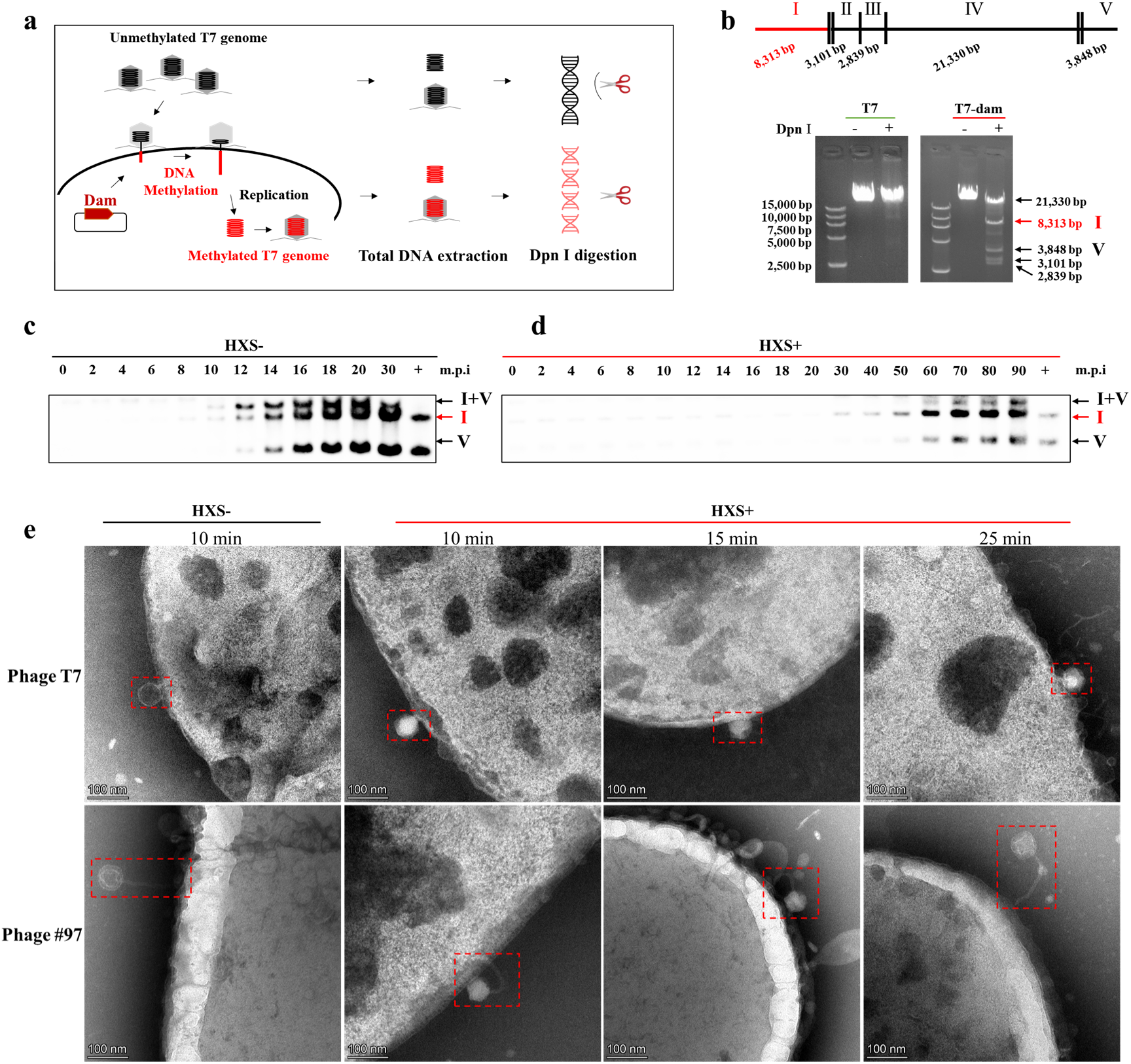
The HXS system disrupts phage genome entry. **a**, Schematic diagram showing that *Dpn*I specifically cleaves methylated, but not unmethylated, T7 DNA. *E. coli* BL21 cells overexpressing Dam methylase were infected with the unmethylated T7 phage, whose progeny genomic DNAs were methylated and can be cleaved by *Dpn*I. **b**, *Dpn*I digests the T7 genomic DNA prepared by infection of regular *E. coli* BL21 cells (T7) or *E. coli* BL21 cells overexpressing Dam methylase (T7-dam). Six *Dpn*I cleavage sites and the size of 5 large fragments (I to V) were shown. The left end fragment (I) was highlighted in red. **c-d**, *Dpn*I digestion assay for measurement of DNA translocation rate. *E. coli* BL21 cells overexpressing Dam methylase were transformed with the plasmid expressing the HXS system (HXS+) or vector control (HXS-), and infected with T7 phage. At different time points after infection, the infection mixtures were collected to isolate T7 DNA, which was then digested with *Dpn*I. After agarose gel electrophoresis, the DNA was transferred to nylon membranes and analyzed by southern blot using the fragment I-specific and fragment V-specific probes. “I+V” denotes a restriction fragment that can be detected by both the I and V probe; the “V” fragment is detected only by probe V. **e**, Transmission electron microscopy images of phages. Empty capsids display a dark core due to the electron-dense staining material, while full capsids show a bright central region.

We found that in the absence of HXS, fragment I was detected at 8 min post infection (mpi) and fragment V at 12 mpi, consistent with ∼10-12 min for complete T7 entry (Fig. 2c) ^33^. A larger head-to-tail (I+V) band appeared by 12 mpi, indicating the onset of concatemeric replication ^34^. Strikingly, in the presence of the HXS system, fragment I and fragment V were delayed to ∼30 mpi and ∼50 mpi, respectively (Fig. 2d), demonstrating a substantial retardation of genome entry. Consistently, the HXS-escape T7 mutant can inject its genomic DNA into *E. coli* BL21 cells expressing HXS at the similar efficiency (Supplementary Fig. 4).

To visualize genome entry directly, we performed time-resolved transmission electron microscopy (TEM), where DNA-emptied capsids stain dark, whereas DNA-filled capsids appear light ^35^. Without HXS, most T7 capsids were dark by 10 min post infection (mpi), consistent with completed genome ejection, whereas in HXS-expressing cells, capsids remained largely light even at 25 mpi (Fig. 2e), in agreement with the delay in DNA entry observed in the *Dpn*I assay. Because T7 completes genome delivery via a transcription-dependent mechanism, in which host σ⁷⁰ RNA polymerase initiates from early promoters A1, A2, and A3 before takeover by the viral polymerase gp1 ^36^, we asked whether HXS might act primarily by blocking these early transcriptional events. Deletion of the σ⁷⁰-dependent A2 and A3 promoters from the T7 genome did not relieve HXS-mediated inhibition in plaque assays (Supplementary Fig. 5a-b). Moreover, when cells carrying an A1 promoter driven GFP reporter plasmid were infected with either wild-type T7 or the ΔA2A3 mutant, HXS neither reduced its restriction efficiency nor detectably altered A1-driven GFP expression (Supplementary Fig. 5c-d). Together with the entry assays above, these data indicate that HXS does not simply inhibit the transcriptional program required for T7 DNA translocation, but instead targets the genome-entry step itself.

To test whether entry inhibition generalizes beyond T7, we randomly picked up a second coliphage from the family Christensenvirus and examined the DNA entry. TEM at matched time points showed the same pattern: without HXS, capsids appeared dark by 10 mpi, whereas with HXS, capsids remained light at 25 mpi (Fig. 2e). Thus, HXS-mediated delay of DNA entry is observed across distinct coliphage lineages, reinforcing the conclusion that HXS targets the genome-delivery step of infection. Together, these results indicate that the HXS interferes with phage DNA entry.

### The periplasmic localization of cleaved HxsA protein is indispensable for the antiphage activity of the HXS system

Since the HXS system interferes with phage DNA entry, we hypothesized that some of the four HXS proteins might localize to the bacterial wall. Sequence analysis showed that only the HxsA protein possesses a signal peptide (residues 1-22) and a peptidoglycan-binding domain (Fig. 3a and Supplementary Fig. 6a). Replacement of the signal peptide with a hexa-histidine tag (HxsA_Rep-SignalP_) or mutations on peptidoglycan-binding domain abolished the antiphage activity (Fig. 3b and Supplementary Fig. 6b-c), indicating that the HxsA protein may locate in periplasm and act as an effector of the HXS system.

**Fig. 3.**
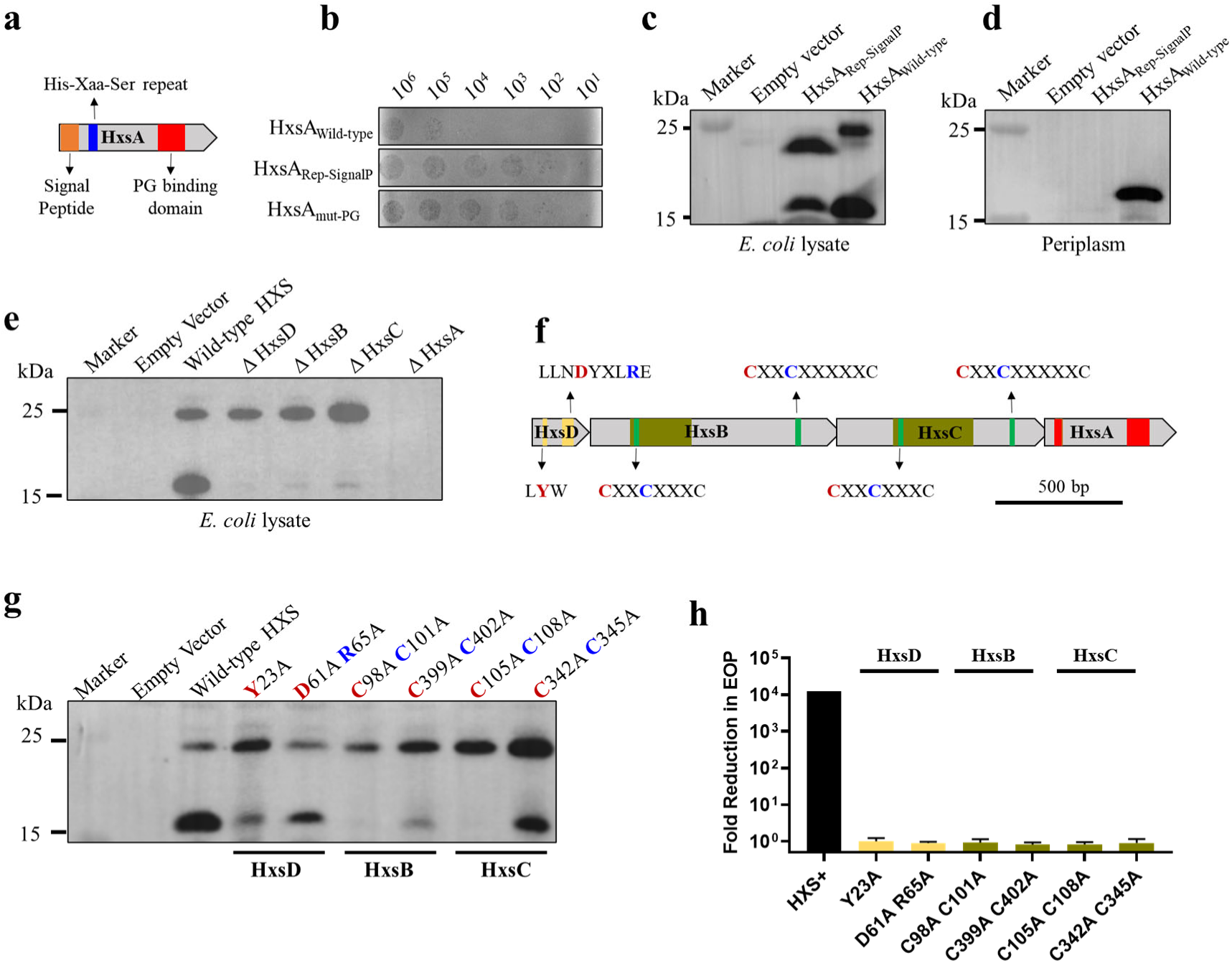
The periplasmic localization of cleaved HxsA protein is indispensable for the antiphage activity of the HXS system. **a**, Schematic diagram showing the functional domains of HxsA protein. **b**, Plaque assay of T7 phage infecting *E. coli* cells expressing the HXS systems with wild-type or mutated HxsA proteins. The HxsA_Rep-SignalP_ have 6˟His tag replacing the signal peptide of the HxsA protein, whereas HxsA_mut-PG_ contains targeted mutations in the peptidoglycan-binding region. **c-d**, Subcellular localization analysis of wild-type and mutant HxsA proteins by western blot. Whole-cell lysates (**c**) and periplasmic fractions (**d**) of *E. coli* were prepared as described in the Materials and Methods and analyzed by western blot using HxsA-specific polyclonal antibodies. **e-g**, Western blot analysis to determine the essential genes and the critical amino acid residues for the cleavage of the HxsA proteins. **e**, Western blot analysis of the cleavage efficiency of the HxsA protein in the single gene deletion HXS systems. **f**, Schematic diagram showing the conserved motifs of the HXS system. The mutated amino acid residues were highlighted in blue or red. **g**, Western blot analysis of the cleavage efficiency of the HxsA protein with different mutations. The wild-type HXS system was used as a control. **h**, The sites mutant analysis of HXS system. Wild-type and the mutated HXS systems were transformed into in *E. coli* BL21, which were infected with T7 phage. Empty vector was used as a control.

Periplasmic proteins of *E. coli* were prepared as described previously ^37,38^ and analyzed by western blot using home-made HxsA-specific polyclonal antibodies (Supplementary Fig. 7). Although both wild-type HxsA and HxsA_Rep-SignalP_ were detected in whole-cell lysate, only wild-type HxsA was recovered from the periplasm (Fig. 3c-d). Notably, HxsA_Rep-SignalP_ was still cleaved, albeit with reduced efficiency, indicating that the observed proteolysis is not simply canonical signal-peptide removal. Indeed, single-gene deletions showed that each of the other Hxs proteins is necessary for HxsA cleavage (Fig. 3e), and fusing the predicted HxsA signal peptide to GFP did not direct GFP to the periplasm (Supplementary Fig. 8). Together, these results indicated that the signal peptide alone is insufficient for secretion and does not behave as a canonical *E. coli* signal peptide.

Since all the HxsB, HxsC, and HxsD lack a signal peptide, they might participate the cleavage of HxsA in the cytoplasm. To experimentally assess their localization, we generated polyclonal antisera against overexpressed HxsB, HxsC and HxsD (Supplementary Fig. 9), but endogenous levels in cells expressing the native HXS system were below the detection limit of Western blotting, so we cannot yet definitively assign their precise subcellular localization. However, single gene deletion showed that all these proteins are necessary for the HxsA modification (Fig. 3e). HxsB and HxsC proteins are characterized by the radical SAM enzyme with two 4Fe-4S cluster domains (Fig. 3f). Double mutation of the first 4Fe-4S cluster domain of HxsB (C98AC101A) or HxsC (C105AC108A) completely inhibited the HxsA cleavage (Fig. 3f-g), while double mutation of the second 4Fe-4S cluster domain of HxsB (C399AC402A) or HxsC (C342AC345A) decreased the cleavage efficiency. We did not examine single-residue substitutions, which may retain partial cofactor binding and residual activity. HxsD is a small protein belongs to TIGR03976 superfamily, many members of which contain conserved motifs LYW and LLNDYxLRE. Targeted substitutions in HxsD revealed that the LYW motif (Y23A) and the LLNDYxLRE motif (D61A/R65A) also reduce the cleavage efficiency (Fig. 3f-g). In the D61A/R65A mutant, the cleaved HxsA fragment is indeed still abundant, but the fraction of cleaved versus uncleaved HxsA is clearly reduced compared with the wild-type. This pattern indicates that LLNDYxLRE motif partially impairs HxsA processing, with a milder effect than the other mutations. In addition, mutating either of these key motifs of HxsB, HxsC, and HxsD abolished the antiphage activity (Fig. 3h).

Together, these results indicated that the HxsB, HxsC, and HxsD work together to cleave the HxsA, which is necessary for its antiphage activity.

### The rSAM enzyme-mediated maturation of HxsA establishes a periplasmic barrier to phage DNA entry

To further characterize the cleaved HxsA, the band corresponding to the cleaved HxsA was excised from the SDS-PAGE gel (Supplementary Fig. 10), and the proteins were extracted and digested with trypsin and chymotrypsin, respectively, for mass spectrometric analysis. LC-MS/MS analysis of the cleaved HxsA protein identified eight tryptic and nine chymotryptic peptides, collectively covering nearly all residues 48-214 of HxsA (Fig. 4a). Two fragments, residues 23-47 and 114-139 (highlighted in cyan in Fig. 4a), were not detected. Mutation analysis showed that residues 23-47, but not the residues114-139, are required for antiphage activity (Fig. 4b). Because residues 23-47 harbor four chymotryptic sites, this region was likely fragmented into peptides below the MS detection limit. To determine whether the periplasmic localization of cleaved HxsA alone is sufficient for defense, we expressed four N-terminal truncations starting at each chymotryptic site residues 23-47 (Fig. 4a), each fused to either a Tat or Sec signal peptide (eight constructs; Supplementary Fig. 11a). Western blots confirmed that all eight variants were transported to the periplasm, yet none conferred antiphage activity (Supplementary Fig. 11b-c). Thus, simply placing cleaved HxsA in the periplasm is insufficient; HxsB, HxsC, and HxsD are required beyond cleavage.

**Fig. 4.**
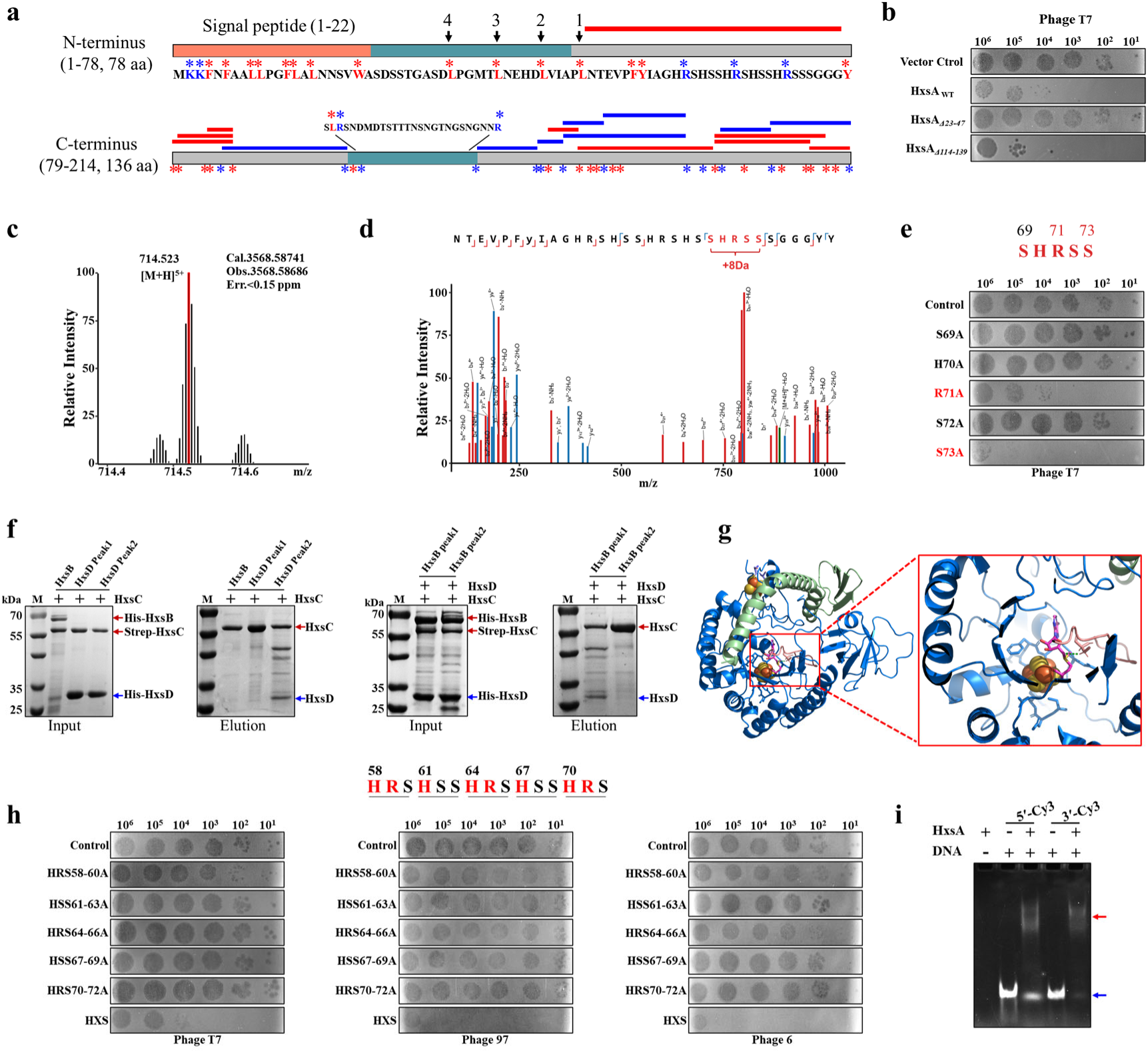
The rSAM-mediated maturation of HxsA establishes a periplasmic barrier to phage DNA entry. **a**, Schematic diagram showing the results of the spectrometric analysis of the cleaved HxsA protein. The band corresponding to the cleaved HxsA protein was excised from the SDS-PAGE gel. The HxsA proteins were subtracted, digested with trypsin and chymotrypsin respectively, and subjected to mass spectrometric analysis. Red and blue stars indicate the cleavage sites of chymotrypsin and trypsin, respectively. The red and blue lines indicate the identified chymotrypsin-digested and trypsin-digested peptides, respectively. The residues 23-47 and 114-139 that was not detected was highlighted in cyan. **b**, Plaque assay analysis of the effect of the residues 23-47 and 114-139 on the antiphage activity of HXS. **c-d**, Identification of the covalently modified peptides by mass spectrometry. **c**, High-resolution mass spectrum of the precursor ion at m/z 714.5, corresponding to the [M+H]⁵⁺ charge state. **d**, Tandem mass spectrum (MS/MS) of modified peptide. The fragmentation patterns that generate the observed b and y ions are illustrated along the peptide sequence shown on top of the spectrum. b ions are colored red, and y ions are colored blue. **e**, Plaque assay analysis of the effect of a single-residue mutation within the SHRSS motif on the antiphage activity of HXS. **f**, Pull-down assays show the interaction between HxsC and HxsD. The presence of HxsB, HxsC and HxsD was analyzed by SDS-PAGE. **g**, Structural model of the HxsC-HxsD complex engaging an HxsA-derived peptide. A docking model showing the interaction between HxsC (blue) and HxsD (green), with a short peptide (pink) from HxsA containing the SHRSS motif positioned near the radical SAM active site of HxsC. The [4Fe-4S] cluster is shown as spheres, and SAM is depicted in stick (magenta). **h**, Plaque assay analysis of the effect of a single HxS repeat on the antiphage activity of HXS. Residues with positive charges are highlighted in red. **i**, Periplasmic HxsA isolated from bacteria expressing HXS binds DNA. The shifted band indicates DNA-protein complexes (red arrow), whereas free DNA migrates faster (blue arrow).

Notably, liquid chromatography-high-resolution mass spectrometry (LC-HRMS) detected a peptide matching HxsA (48-79) with a +8.0 Da mass shift relative to the unmodified species (hereafter denoted HxsA (48-79) [+8]) in the precursor spectrum. High-resolution MS/MS of HxsA (48-79) [+8] yielded diagnostic fragment ladders that unambiguously localized the +8 Da modification to the SHRSS motif within this segment (Fig. 4c-d). The observed isotopic pattern and low-ppm mass error were consistent with a discrete +C/–4H–type alteration rather than adduct formation or oxidation. Site-directed alanine scanning of the SHRSS motif revealed that substituting either the N-terminal Ser (S to A), His (H to A), or C-terminal Ser (S to A) abolished HXS-mediated protection (Fig. 4e), indicating that the +8 Da modification mapped to SHRSS is essential for antiphage function (Fig. 4d-e). However, because mutation of SHRSS or deletion of *hxsB*, *hxsC*, or *hxsD* severely impaired HxsA processing and periplasmic accumulation, we could not recover HxsA from these backgrounds for LC-MS/MS; accordingly, we were unable to directly assess the +8 Da modification state of HxsA in the Δ*hxsB*/Δ*hxsC*/Δ*hxsD* or active-site mutant.

Because HxsB and HxsC each contain an rSAM domain, a hallmark of enzymes involved in peptide and protein modification ^20^, we hypothesized that they catalyze a covalent maturation of HxsA, consistent with our genetic data showing that active-site substitutions in either protein abolish HxsA cleavage and defense (Fig. 3g-h). To probe how this maturation machinery is organized, we mapped physical interactions among the HXS components. Pull-down assays revealed a HxsC-HxsD complex, whereas no stable association between HxsB and HxsC was detected (Fig. 4f and Supplementary Fig. 12a-c), indicating that HxsC engages HxsD to form the core processing module. Consistently, AlphaFold 3 modelling predicted an HxsC-HxsD heterocomplex with lower free energy than alternative assemblies (Supplementary Fig. 13a-c). Docking the HxsA segment encompassing the SHRSS motif into this model positioned the histidine and terminal serine side chains near the unique-iron/SAM pocket in HxsC, a geometry compatible with the observed +8 Da mass shift on SHRSS and with a site-specific covalent modification at these residues (Fig. 4g). These structural predictions support a model in which the HxsC-HxsD core complex installs a covalent modification on SHRSS motif, although further studies are required to define the precise mechanism.

The breadth of HXS activity suggests that it targets a phage-agnostic step rather than specific structural proteins. Consistent with this, no detectable interaction was observed between periplasmic HxsA and the T7 ejectosome proteins gp15/gp16 in pull-down assays (Supplementary Fig. 14a-d). HxsA carries five tandem HxS repeats enriched in basic residues, including histidine that provides pH-dependent positive charge capable of electrostatic interactions with DNA. We therefore hypothesized that electrostatic engagement of incoming DNA underlies inhibition of genome entry. Functional scanning supported this model: triple-alanine replacement of any single HxS repeat (HxS to AAA) abolished HXS-mediated protection (Fig. 4h), indicating that the repeats act as a multivalent functional array. Moreover, fluorescent DNA-binding assays with periplasmic HxsA showed robust DNA binding (Fig. 4i). Together, these data argue that HxsA does not rely on phage-specific protein recognition; instead, its positively charged, repeated HxS motifs likely capture or sequester the translocating genomic DNA within the periplasm, providing a mechanistic basis for the system’s broad antiphage spectrum.

Taken together, these results suggest that HxsB, HxsC, and HxsD act in the cytoplasm to maturate HxsA, including installing a site-specific +8 Da modification within the last HxS repeat and promoting N-terminal processing. The mature HxsA is exported to the periplasm, where its peptidoglycan-binding domain anchors it to the peptidoglycan layer. Its five basic-residue-enriched HxS repeats, including the modified motif, most likely engage the incoming phage genomic DNA by electrostatic interactions, thereby impeding DNA entry and conferring broad antiphage defense (Fig. 5a).

**Fig. 5.**
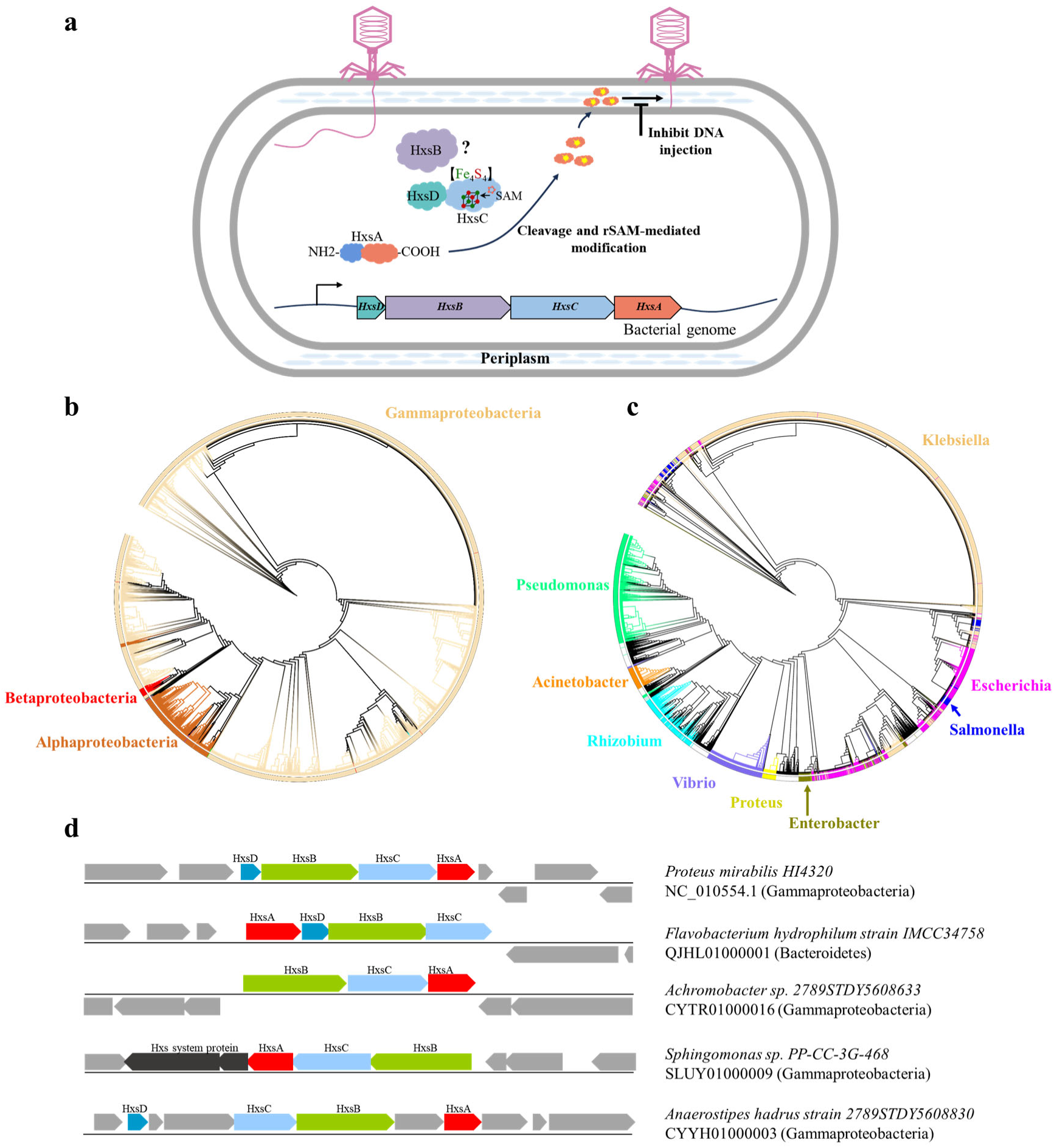
Proposed model and distribution of the HXS system. **a**, Proposed model of the HXS system. HxsC, a radical S-adenosylmethionine enzyme containing a [4Fe-4S] cluster, forms a complex with HxsD and, together with HxsB, processes HxsA in the cytoplasm, including N-terminal cleavage and installation of a site-specific modification on the SHRSS motif. The matured HxsA effector is then exported to the periplasm, where its peptidoglycan-binding domain anchors it at the cell wall and its five HxS repeats likely interfere with phage DNA entry. **b-c**, Clustering of the identified 10,369 HxsA proteins at the level of bacterial phylum (**b**) and genus (**c**). Different bacterial genera are shown in different colors. **d**, Gene structure of the HXS systems.

### The distribution and gene structure of the HXS system

Since the HxsA is the effector protein of the HXS system, we determined the distribution of the HXS system by evolutionary analysis of HxsA proteins across all bacteria. Our results showed that 99.7% of the HXS system was found in the phylum *Pseudomonadota*, whereas there was only a sporadic distribution in other bacterial phyla. *Gammaproteobacteria* accounted for 91.3% of the HXS system-containing bacteria, which was much more than the other bacterial phyla (Fig. 5b).

A species breakdown of HXS-containing bacteria revealed that *Klebsiella* is the most prevalent species of HXS-containing bacteria, and 27.2% of the 16,000 *Klebsiella* genomes in the NCBI database (through January 2022) contain the HXS system (Fig. 5c). This suggests that the HXS system may be an important defense system within *Klebsiella*. In addition, this system is also abundantly distributed in *Escherichia* and *Pseudomonas* (Fig. 5c). Evolutionary analysis showed that the HXS systems of *Pseudomonas*, *Acinetobacter*, *Rhizobium*, *Vibrio*, *Proteus*, and *Enterobacter* were well separated from other bacteria. However, the HXS systems of *Escherichia*, *Salmonella* and *Klebsiella* are distributed in different branches and cross each other (Fig. 5c), indicating that the HXS system may be transmitted in these bacterial species.

Gene structure analysis showed that the composition of proteins within the HXS system is relatively conservative, of which more than 99.8% is composed of four proteins, HxsA, HxsB, HxsC, and HxsD (Fig. 5d). HxsB and HxsC are always close together, while HxsA and HxsD are sometimes separated by other proteins. In a few cases the HxsD protein is missing. In addition, 99.6% of HXS systems are organized in the order HxsD-HxsB-HxsC-HxsA, and only a small number of systems have different gene organizations (Fig. 5d).

## DISCUSSION

Despite the extensive catalog of bacterial antiphage systems have been identified, surprisingly, no system that specifically targets the phage genomic DNA entry phase has been reported until this study. Our discovery of the HXS system fills this gap by introducing a mechanism that uniquely disrupts phage DNA entry, thereby adding a critical layer to our understanding of bacterial immune defense. This innovative mechanism not only prevents the progression of the infection, but also conserves energy by minimizing the metabolic costs typically associated with more destructive pathways, such as abortive infection systems. Such a strategy underscores a novel layer of bacterial immune defense.

Distinct from phage-encoded superinfection exclusion (Sie) systems ^19^ such as the Imm and Sp systems of T4 phage, which act as narrow-spectrum self-protection mechanisms against closely related phages, the HXS is an intrinsic bacterial defense system that provides broad and potent antiphage activity (Fig. 1). Whereas T4 Sie systems block DNA entry during superinfection to favor its replication, HXS disrupts phage DNA entry to protect the bacterial cell itself against a wide range of coliphages. This fundamental difference underlines the role of the HXS system in the ecological and evolutionary dynamics of microbial communities, suggesting a key adaptation for bacterial competitiveness and survival.

The HxsA protein contains five HXS repeats, which had been identified in bacterial proteins for over a decade, but their function has remained elusive ^21^. Radical SAM enzymes catalyze diverse radical transformations, including the maturation and modification of peptides and proteins ^39,40^. SAM-linked defense systems also exist: BREX uses SAM in host defense, while certain phages encode a counter-defense SAM lyase that depletes SAM ^41^, and type III CRISPR employs SAM-AMP as a signaling nucleotide ^42^. However, these mechanisms do not entail rSAM enzyme-catalysed covalent modification of a protein effector. Our work assigns a defense role to an HxS-repeat protein and links it to rSAM enzyme-guided maturation: HxsB, HxsC and HxsD are required for N-terminal processing of HxsA and for a site-specific +8 Da modification within an HxS repeat. The mature HxsA is then exported to the periplasm, where it executes its antiphage function.

Many bacterial systems recognize one or a few phage proteins or sense a single intracellular event during replication, which constrains the spectrum of phages defended. By contrast, HXS shows a broad antiphage spectrum across diverse coliphage lineages. One parsimonious explanation is its target choice: rather than phage-specific protein cues, HXS most likely engages the incoming DNA itself. In our model, mature HxsA resides in the periplasm and is retained at the peptidoglycan layer via its peptidoglycan-binding module. Its five tandem HxS repeats enriched in basic amino acids, including the site-specifically modified motif, most likely electrostatically attract and perturb the translocating genome, thereby impeding entry. This DNA-centric mechanism provides a basis for HXS’s broad-spectrum defense while also explaining rare escapes in which entry dynamics or periplasm-facing interfaces differ.

Although HXS acts at the DNA-entry step and restricts most coliphages tested, we have not yet examined its impact on plasmid transduction or on other mobile genetic elements such as phage satellites. Notably, the classical generalized transducing phage P1 is not restricted by HXS under our experimental conditions and behaves as an escape phage in our assays (Fig. 1a). This informative exception suggests that differences in DNA-entry mechanisms or periplasm-facing interfaces may allow certain phages to evade periplasmic interception by HxsA. Phage P1 therefore provides a promising starting point for future transduction-focused studies aimed at dissecting how HXS shapes the spread of plasmids and other mobile genetic elements.

Several technical limitations currently preclude a complete biochemical and structural reconstruction of the HxsB/HxsC/HxsD maturation module. When expressed from the endogenous promoter, HxsB, HxsC and HxsD are below the detection limit of Western blotting (Supplementary Fig. 9), making it currently impossible to purify sufficient native complex for in vitro reconstitution. Although HxsB, HxsC and HxsD can be individually overexpressed, overexpression of HxsA alone yields predominantly insoluble inclusion bodies, and we have so far been unable to reconstitute rSAM enzyme-dependent modification of HxsA *in vitro* from recombinant components. Consistent with this, mutations in *hxsB*, *hxsC* or *hxsD* abolish processing and periplasmic accumulation of HxsA, preventing isolation of fully matured HxsA from these backgrounds. These constraints highlight the likely importance of the native cellular environment and/or additional host factors for efficient HxsA maturation and export, and underscore that the precise radical chemistry underlying the +8 Da modification and the detailed structural organization of the maturation complex remain to be elucidated.

In conclusion, this work identified, to our knowledge, the first rSAM enzyme-dependent covalent modification of a protein effector implicated in bacterial antiphage defense, and proposes a periplasmic interception strategy at the DNA-entry step. By acting upstream of genome replication, HXS broadens the mechanistic landscape of bacterial immunity. In addition, the broad-spectrum antiphage activity of HXS points to opportunities for engineering phage-resistant strains for biotechnology.

## METHODS

### Bacterial strains, phages, and plasmids

*E. coli* DH5α was used for plasmid construction, and *E. coli* B834 for engineering T7 phage mutants. *E. coli* MG1655 served as the host for initial antiphage activity screens, and *E. coli* BL21 was used for mechanistic assays of the HXS defense system. The Zorya type II and RT-TIR defense systems were cloned in previous studies ^16,43^. The HXS, RT-Abi-P2, and AVAST type 2 systems were PCR-amplified from genomic DNA of *E. coli* strains MEM (GenBank accession CPO12378), RHB26-C20 (GenBank accession CP057447), and EC6030 (GenBank accession CP183338), respectively, and assembled into pSEC1 by standard methods. The resulting plasmids were individually transformed into *E. coli* MG1655 and used for antiphage assays. The HXS systems with different mutations were constructed by one-step cloning (Supplementary Table 3). T7 phage (GenBank Accession V01146.1) was maintained in our laboratory stocks. The panel of 109 phages isolated on *E. coli* MG1655 has been described previously ^43^ and used for antiphage assays. Genome sequences for these 55 isolated phages were sequenced and deposited in NCBI (GenBank Accession numbers were listed in Supplementary Table 1).

### Bioinformatic analysis

A total of 2,289 complete *E. coli* genome sequences available by October 2020 were downloaded from NCBI GenBank. The genes were annotated using Prodigal version 2.6.3 software, and the resulting protein sequences were functionally annotated and assigned to cluster of orthologous groups (COG) categories using Eggnog-mapper v2.1.6. The homologs of the eukaryotic viperin protein (rSAM (COG 0535)) were searched within the annotated COG category. All COG0535 proteins were then analyzed for nearby proteins to determine the operon structure. The conserved domains of the HXS system were identified using Motif Search ^44^. The signal peptides of the HxsA protein were predicted using SignalP 6.0 ^45^.

### Phage challenge assay

Phage challenge assays were performed as described previously ^43^. Briefly, empty vector or pSEC1 constructs expressing individual defense systems were transformed into *E. coli* MG1655 cells. Transformants were grown with the appropriate antibiotics to mid-log phase (typically OD₆₀₀ ≈ 0.3-0.4). For each assay, 300 μL of culture was mixed with 8 mL of top agar containing the same antibiotics and poured evenly onto LB agar plates. Each of 113 phages (111 phages mentioned above plus T4 and P1 phages) was tenfold serially diluted individually, and 1 μL of each dilution was spotted onto the bacterial lawn. Plates were incubated overnight at 37 °C, and plaque formation was recorded. Antiphage efficacy was expressed as the endpoint-dilution ratio: the highest dilution that produced ≥1 plaque on the empty-vector lawn divided by the highest dilution that produced ≥1 plaque on the defense-system lawn. Unless stated otherwise, data represent ≥3 biological replicates.

### Phage-infection dynamics in liquid medium

Overnight culture of *E. coli* BL21 (DE3) expressing HXS defense system (or empty vector control) was inoculated into fresh LB medium supplemented with 50 µg/mL kanamycin and incubated at 37 ℃ with a shaking speed of 200 rpm. When cultures reached 4×10^7^ cells/mL, 180 µL aliquots were dispensed into microwells of a 100-well plate, and 20 µL of phage suspension was added to achieve multiplicities of infection (MOI) of 5 and 0.05 (two parallel conditions). Plates were instantly placed in an automated growth curve analysis system (Oy Growth Curves Ab Ltd Bioscreen C), and OD_600_ of the culture was recorded every 10 min for 8h at 37 °C with continuous shaking. Bacterial growth curve was plotted using GraphPad Prism software.

### Phage adsorption assay

Overnight cultures of *E. coli* BL21 (DE3) containing the HXS system or the empty control were transferred to fresh LB medium at a ratio of 1:100 and incubated at 37 ℃ with a shaking speed of 200 rpm. When cultures reached 1×10^8^ cells/mL, T7 phage were added to a final concentration of 2×10^3^ PFU/mL, and the infection mixture was incubated at 37 ℃ with a shaking speed of 200 rpm. At 0, 1, 5, 10, and 15 minutes after infection, 1 mL of the infection mixture was collected and centrifuged at 12,000 g for 90 seconds. 100 uL of the supernatant was collected and titrated for the unadsorbed free phage by plaque assay.

### Selection of T7 phage mutants escaping the HXS system

300 μL of an overnight culture of *E. coli* BL21 (DE3) cells expressing the HXS system was mixed with 8 mL of top agar and poured onto LB agar plates. Tenfold serial dilutions of T7 were prepared, and 2.5 µL of each dilution was spotted onto the HXS lawn. Plates were incubated overnight at 37°C. The resulting plaque was collected into an Eppendorf tube containing 1 mL of Pi-Mg buffer. After 1 h incubation at room temperature with gentle vertexing, phage was titrated and used for the next round of selection (serial passage) using the same procedure. Rounds of selection were repeated until T7 formed plaques on the HXS-expressing BL21(DE3) lawn with an efficiency comparable to the empty-vector control lawn. After the final round, escape phage was purified by single-plaque isolation prior to amplification and downstream analyses.

### Genomic sequencing of escaped T7 phage mutants

Escaped phages were propagated as described previously ^43^. Briefly, 1 L of *E. coli* BL21(DE3) cells were infected with the escape phage at a MOI of 0.05. After 5 hours incubation at 37°C, *E. coli* cells were removed by centrifugation at 5,600 g for 20 min, and the supernatant was filtered through a 0.22 μm filter to further remove bacterial debris. The phages in the supernatant were pelleted at 30,000 g for 30 min at 4 °C, then resuspended in Pi-Mg buffer. For genomic DNA extraction, phage resuspensions were treated with DNase I (10 μg/mL) and RNaseA (10 μg/mL) to remove contaminating bacterial DNA. EDTA (5 mM) and SDS (35 mM) were then added to inactivate nuclease and lyse phages for 20 minutes at 65°C. Lysates were then treated with phenol/chloroform/isoamyl alcohol (25:24:1), followed by chloroform/isoamyl alcohol (24:1) to remove proteins. T7 genomic DNAs in the aqueous phase was precipitated with isopropanol, pelleted by centrifugation at 11,000 g, and washed twice with 70% ethanol. The T7 genomic DNA precipitate was dissolved in distilled water and was sent for sequencing. Mutation sites were identified by sequence alignment of wild-type and mutant T7 genome. Primers were designed to amplify the mutation sites, and PCR products were sequenced by sanger sequencing to further confirm the mutations.

### Genome editing of T7 phage

Recombinant T7 phages carrying defined mutations in gene *15* or gene *16* were generated individually using CRISPR-Cas technology as described previously ^46^. Briefly, five CRISPR-Cas12a plasmids, each expressing a single spacer targeting one of the five mutation sites, were constructed. The sequence of spacers is shown in Supplementary Table 4. Donor plasmids containing the desired mutation and flanking homology arms were constructed using standard methods. For each edit, donor plasmid and the corresponding CRISPR-Cas12a plasmid were co-transformed into *E. coli* B834 cells. 100 μL of Pi-Mg buffer containing different number of wild-type T7 phages were mixed with 300 μL of the freshly cultured *E. coli* B834 cells, poured onto LB agar plates, and incubated overnight at 37°C to generate the T7 phage mutants. *E. coli* B834 cells containing only the CRISPR-Cas12a plasmid were used as controls. Individual plaques obtained on the co-transformed strain were picked and plaque-purified, and the targeted loci were PCR-amplified and verified by Sanger sequencing to confirm the intended mutations.

### Preparation of T7 phage with methylated genomic DNA

The *dam* gene was PCR-amplified from the *E. coli* strain MG1655 and cloned into pET-22b expression vector to generate pET-Dam plasmid, which was then transformed into *E. coli* BL21. Overnight cultured *E. coli* BL21 cells were inoculated into fresh LB medium at a 1:100 dilution and grown at 37 °C with shaking. When cultures reached OD_600_=0.6, expression of Dam was induced with IPTG (1 mM) for 2 hours. *E. coli* BL21 cells were then infected with T7 phage at a MOI of 0.4. After 3 hours incubation at 37 °C, the infection mixture was collected and phages were prepared as described above. Methylation of T7 genomic DNA was confirmed by *Dpn*I digestion, which specifically cleaves the Dam-methylated DNA. T7 phage prepared by infecting *E. coli* BL21 cells with pET-22b empty vector was used as a control.

### Southern blot

*E. coli* BL21 cells containing pET22b-Dam were transformed with the HXS system-expressing plasmid or empty vector. Transformants were grown in LB medium at 37 °C until the OD_600_ reached 0.6, then infected with T7 phage at a MOI of 0.4. 100 μL of the infection mixture were collected every 2 min for the first 20 min, and every 10 min thereafter up to 90 min. Each sample was immediately mixed with an equal amount of phenol/alcohol solution. After centrifugation at 12,000 rpm for 2 min, the pellet was resuspended with Tris-HCl buffer containing proteinase K (100 μg/uL), followed by incubation at 37 °C for 1 h. T7 phage genomic DNAs were extracted using the phenol-chloroform-isoamyl alcohol as described above and treated with *Dpn*I (1 μL/μg) at 37 °C for 1 h to digest methylated DNA. The DNA samples were then analyzed by electrophoresis on 0.8% agarose gels and transferred to nylon membranes using the capillary transfer method with an alkaline transfer solution (400 mM NaOH, 1 M NaCl). Membrane was equilibrated in 6×SSC buffer (0.9 M NaCl, 0.09 M sodium citrate tribasic dehydrate) for 2 min and pre-hybridized in hybridization buffer (6×SSC, 5×Denhardt solution, 0.5% SDS and 1 mg/ml salmon sperm DNA) for 2-4 h at 42 ℃. Head and tail fragments of the T7 phage genome were individually amplified by PCR (Supplementary Table 5), biotin-labelled using a Biotin Random Prime DNA Labeling Kit (Beyotime, China), and used as probes for Southern blot. Membranes were then hybridized overnight with biotin-labelled DNA probes at 42 ℃ and developed with the chemiluminescence detection kit (Beyotime, China) according to the manufacturer’s instructions.

### Expression and purification of HxsA, HxsB, HxsC, and HxsD protein

Recombinant HxsA, HxsB, HxsC, and HxsD proteins each with a hexa-histidine tag were purified by nickel-based affinity chromatography. Briefly, the *HxsA, HxsB, HxsC, and HxsD* genes were cloned into the pET32a vector. HxsA proteins were induced with 1 mM IPTG at 28°C for 5 hours upon reaching an OD_600_ of 0.6-0.8, whereas HxsB, HxsC and HxsD were induced at 16 °C overnight. A Strep-tagged HxsC protein was generated for pull-down assays and expressed in *E. coli* under the same induction conditions. For His-tagged proteins, *E. coli* cells were harvested, resuspended in binding buffer (20 mM Tris-HCl pH 8.0, 100 mM NaCl, 10 mM imidazole, and 5 mg/mL DNase I), and disrupted by high-pressure cell disruption. Cell debris was removed by high-speed centrifugation, and proteins were then purified using a HisTrap column and analyzed by SDS-PAGE. For Strep-tagged HxsC, cells expressing Strep-HxsC were resuspended in equilibration buffer (100 mM Tris pH 8.0, 150 mM NaCl), lysed and purified using Strep-Tactin affinity resin. Eluted proteins were further purified by size-exclusion chromatography. All purified proteins were analyzed by SDS-PAGE, and concentrations were estimated using a BSA standard.

### Preparation of the HxsA-, HxsB-, HxsC-, and HxsD-specific polyclonal antibodies

Six-week-old female BALB/c mice were purchased from Laboratory Animal Center of Huazhong Agricultural University (Wuhan, China). Animal experiments were performed according to the protocols approved by the Animal Protection and Ethics Committee of Huazhong Agricultural University (HZAUMO-2023-0094), Hubei, China. The purified HxsA, HxsB, HxsC, and HxsD proteins were emulsified with complete Freund’s adjuvant for the primary immunization and with incomplete Freund’s adjuvant for booster immunizations, and administered subcutaneously to mice (n=3, 100 μg of each protein per dose) at weeks 0 and 2. Mice (n=3) immunized with PBS were used as controls. Serum samples were collected 14 days after the last immunization, and antigen-specific antibody titers were determined by ELISA. Sera were aliquoted and stored at -80°C.

### Antibody detection in sera

Serum samples (n = 3) were collected 2 weeks after the second immunization. ELISA plates were coated overnight at 4 °C with 200 ng/well of purified His-tag HxsB, HxsC, or HxsD protein. Plates were washed five times with PBS containing 0.05% Tween 20 (PBST) and blocked with 3% BSA in PBST for 1 h at 37 °C. Serum samples were serially diluted in 1% BSA in PBST and incubated on the coated plates at 37 °C for 1 h. Following five washes with PBST, peroxidase-conjugated goat anti-mouse IgG was added and incubated for 1 h at 37 °C. Plates were washed five times with PBS-T, and bound antibodies were detected using 3,3′,5,5′-tetramethylbenzidine (TMB) substrate. The reaction was stopped with 2 M H_2_SO_4_, and absorbance was measured at 450 nm using a microplate reader.

### Western blot

*E. coli* BL21 cells containing the wild-type or mutant HXS systems were cultured overnight, collected by centrifugation, and resuspended in SDS-PAGE loading buffer. Proteins were separated by SDS-PAGE and transferred to 0.22-μm polyvinylidene fluoride (PVDF) membranes. After blocking with 5% skim milk in TBST at 37°C for 2 h, the membranes were incubated with primary antibodies (anti-HxsA, anti-HxsB, anti-HxsC, or anti-HxsD; 1:8,000 dilution in TBST) overnight at 4°C. Membranes were then washed with TBST and incubated with horseradish peroxidase (HRP)-conjugated goat anti-mouse IgG secondary antibodies (1:8,000 dilution in TBST) for 1 h at room temperature. Protein bands were visualized using ECL chemiluminescence (Bio-Rad).

### Preparation of the bacterial cytoplasmic and periplasmic proteins

Cytoplasmic and periplasmic proteins of *E. coli* cells were separated as previously described ^37,38^. Briefly, *E. coli* expressing the HXS system or its mutants were diluted 1:100 in fresh LB medium supplemented with kanamycin and grown overnight at 37 °C with shaking. Cells were collected by centrifugation at 4,816 g for 15 min and resuspended in PBS (pH 7.5). After centrifugation at 20,817g for 3 min, the supernatant was discarded. The cell pellet was gently resuspended in Buffer I (100 mM Tris-HCl pH 8.0, 500 mM sucrose, 0.5 mM EDTA) and incubated on ice for 5 min, followed by centrifugation at 20,817g for 3 min at 4 ℃. The supernatant was discarded, and the pellet was gently resuspended in 1 mM Mg buffer and incubated on ice for 2 min. After centrifugation at 20,817g for 3 min at 4 ℃, the supernatant (periplasmic fraction) was collected. The cell pellet was washed twice with Buffer II (50 mM Tris-HCl pH 8.0, 250 mM sucrose, 10 mM MgSO_4_), and then resuspended in Buffer III (50 mM Tris-HCl pH 8.0, 2.5 mM EDTA). Cells were sonicated on ice and centrifuged at 6,708 g for 10 min at 4 °C to remove cell debris. This process was repeated until no visible bacterial cells or debris remained. The supernatant was subjected to ultracentrifugation at 108,726g for 20 min to separate the cytoplasmic proteins (supernatant) from the membrane fractions (pellet).

### Expression and purification of gp15 and gp16 proteins

The gp15 and gp16 proteins were produced and purified as previously described^28,47^. T7 phage *15* and *16* genes were PCR-amplified from T7 genomic DNA and cloned into the pET-16b expression vector, retaining the native N-terminal His-tag. The resulting plasmids (pET16b-gp15 and pET16b-gp16) was individually transformed into *E. coli* BL21 (DE3) competent cells. Single colonies were cultured overnight in LB medium containing 100 μg/mL ampicillin. Ten mL starter cultures were inoculated into 1 L LB broth supplemented with ampicillin and grown at 37 ℃ with shaking. Production of gp15 and gp16 proteins was induced with 0.5 mM IPTG at 37 ℃ for 2 hours when cultures reached an OD₆₀₀ of 0.6-0.8. Cells were harvested by centrifugation at 4,300g for 15 min at 4 ℃ and resuspended in lysis buffer (50 mM HEPES pH 8.0, 300 mM NaCl, 100 mM NDSB-201, 10% glycerol, and 1 mM PMSF). Cell pellets were lysed at 4 ℃ using a high-pressure cell disruptor, and cell debris was removed by centrifugation at 35,000 g for 20 min at 4°C. The supernatant containing the recombinant proteins was incubated with Ni-NTA resin, washed with 50 mL wash buffer (50 mM HEPES pH 8.0, 300 mM NaCl, 20 mM NDSB-201, 10 mM imidazole, 5% glycerol, and 1 mM PMSF), and eluted with elution buffer (20 mM HEPES pH 8.0, 200 mM NaCl, and 400 mM imidazole). Proteins were further purified by size-exclusion chromatography (Superdex200 10/300 column, GE Healthcare Life Sciences) using gel filtration buffer (50 mM HEPES pH 8.0, 100 mM sodium citrate, and 10 mM MgCl_2_). Peak fractions were collected, and the purity of gp15 and gp16 proteins was analyzed by SDS-PAGE.

### Interaction assay of gp15 or gp16 proteins with cleaved HxsA

To determine whether cleaved HxsA associates with gp15 or gp16, periplasmic protein fractions enriched for cleaved HxsA were prepared as described above. Pull-down assays were carried out using His-tag purification resin (Biyuntian Biotech). Briefly, 20 μg of His-tagged gp15 or gp16 proteins were incubated on ice overnight with 90 μL of periplasmic fraction containing cleaved HxsA protein. A total of 150 μL His-NTA resin was packed into a gravity-flow column, washed with pre-chilled water, and equilibrated with gel filtration buffer (50 mM HEPES pH 8.0, 100 mM sodium citrate, and 10 mM MgCl_2_). The protein mixture was then incubated with the equilibrated resin at 4 ℃ for 7 hours. As a negative control, 90 μL of periplasmic fraction (no hig-tagged gp15 or gp16) were incubated with resin under the same conditions. After incubation, the resin was washed with 10 mL wash buffer (50 mM Tris pH 7.5, 300 mM NaCl, and 20 mM imidazole), then resuspended in 100 μL of buffer (50 mM Tris pH 7.5, and 300 mM NaCl). Resin-bound proteins were analyzed by SDS-PAGE followed by immunoblotting using anti-HxsA antibodies (prepared as described above).

### Mass spectrometry analysis of the cleaved HxsA protein

The cleaved HxsA protein was analyzed using liquid chromatography tandem mass spectrometry (LC-MS/MS). Briefly, the *E. coli* BL21(DE3) cells containing the HXS system was cultured overnight at 37°C and collected by centrifugation at 6,708 g for 10 min. The pellets were resuspended in SDS-PAGE loading buffer, and the cleaved HxsA protein was separated on an SDS-PAGE gel and confirmed by western blotting using HxsA-specific antibodies. The band corresponding to cleaved HxsA on the second SDS-PAGE gel was carefully excised, and the HxsA proteins were recovered using electrophoresis. The sample was sent to the Sangon Biotech (Shanghai, China) for LC-MS/MS analysis. Peptides were generated by in-gel digestion with trypsin and chymotrypsin before LC-MS/MS analysis.

Peptide separation was performed by reverse-phase liquid chromatography coupled to tandem mass spectrometry. MS/MS spectra were acquired in a data-dependent acquisition mode using higher-energy collisional dissociation. Raw MS data were processed using Proteome Discoverer (version 2.5, Thermo Fisher Scientific) and searched against the HxsA protein sequence. Carbamidomethylation of cysteine residues (+57.021 Da) was set as a static modification, whereas oxidation of methionine and tyrosine (+15.995 Da), as well as HxsB/C-catalyzed modifications (+12.000 Da or -2.016 Da) on histidine, serine and arginine residues, were set as dynamic modifications. The precursor ion mass tolerance was set to 10 ppm and the fragment ion mass tolerance to 0.02 Da. Peptide-spectrum matches were filtered using a false discovery rate of 1% at the peptide level. Modification site assignment was confirmed by manual inspection of fragment ion spectra.

### Transmission electron microscopy

Overnight-grown *E. coli* Bl21 (DE3) harboring the HXS defense system or an empty vector were subcultured into fresh LB medium to 1×10^8^ cells/mL and infected with purified phages at a multiplicity of infection (MOI) of 50. Samples were collected at 10-, 15-, and 25-min post-infection (empty vector control at 10 min only), chilled on ice, and centrifuged at 12,000g for 3 min at 4 °C to pellet cells with adsorbed phages. Pellets were washed twice and resuspended in pre-chilled, filter-sterilized Pi-Mg buffer. For transmission electron microscopy, 20 μL of the suspension was applied to copper grids for 1 min, blotted, reapplied once, negatively stained twice with 2% phosphotungstic acid, and imaged using a Talos L120C transmission electron microscope at 120 kV.

### DNA-binding assay

The DNA-binding activity of cleaved HxsA was determined using the periplasmic fraction containing cleaved HxsA prepared as described above. 15 μL of the periplasmic fraction was mixed with 3′- and 5′-Cy3-labeled DNA in 20 μL of DNA-binding buffer (200 mM HEPES, 500 mM NaCl, pH 8.0) and incubated at 37 °C for 1 h. The reaction mixtures were run on a 6.5% native polyacrylamide gel in 0.5× TBE on ice for 1.5 h and visualized by chemiluminescence (Bio-Rad).

### Identification of the HxsA cleavage site and signal peptide characterization

The Sec signal peptide MalE or the Tat signal peptide CueO ^48,49^ were inserted upstream of the HxsA cleavage site identified by mass spectrometry, as well as three upstream chymotrypsin cleavage sites. To determine whether the HxsA signal peptide functions as a canonical signal peptide, the *GFP* gene was inserted downstream of the HxsA signal peptide. All constructs were cloned into the arabinose-inducible pBAD vector, and the resulting plasmids were transformed into *E. coli* BL21, respectively. Overnight-grown bacteria were inoculated into fresh medium, cultured to an OD600 of 0.6-0.8, and induced overnight with 0.2% arabinose. Whole-cell lysates and periplasmic fractions were prepared as described above and analyzed by western blotting with an anti-HxsA or anti-GFP antibodies.

### Interaction assays of HxsB, HxsC, and HxsD

To examine interactions among HxsB, HxsC, and HxsD, proteins were prepared as described above and subjected to pull-down assays using Strep-tag affinity resin (Biyuntian Biotech). Briefly, Strep-tagged HxsC (300 pmol) was incubated with HxsB and HxsD at a molar ratio of 1:3 (HxsC:HxsB or HxsC:HxsD) or 1:3:3 (HxsC:HxsB:HxsD) on ice for 2 h. Strep-Tactin Sepharose resin was washed with water and equilibrated with equilibration buffer (100 mM Tris pH 8.0, 150 mM NaCl). The protein mixture was then loaded onto the equilibrated resin, washed with 8 mL equilibration buffer, and eluted with elution buffer (100 mM Tris pH 8.0, 150 mM NaCl, and 2.5 mM desthiobiotin). Resin-bound proteins were analyzed by SDS-PAGE. The structures of HxsB, HxsC, and HxsD proteins were predicted using AlphaFold3^50^, and interface solvation free energies were analyzed with PDBePISA ^51^.

### Assay of HXS effect on T7 phage transcription and translation

The T7 phage σ^70^ promoter A1, ribosome binding site, and intergenic region were cloned into a T vector by Gibson Assembly, and the *GFP* gene was placed under the control of σ^70^ promoter A1 to generate the plasmid P_A1_-GFP. The plasmid was transformed into *E. coli* MG1655 harboring the HXS system or an empty vector (EV). Colonies were inoculated into LB medium containing ampicillin and kanamycin and grown overnight. Plaque assays were performed to assess defense against wild-type T7 phage and a T7 mutant lacking the σ^70^ promoters A2 and A3 (previously constructed in the laboratory). GFP expression was quantified by western blotting using anti-GFP antibodies, with GAPDH as a loading control.

## ACKNOWLEDGEMENTS

This work was supported by the National Natural Science Foundation of China (32570051 and 32503044), Hubei Special Project for Science Development (2024CSA060), Fundamental Research Funds for the Central Universities (2662025DKPY009 and 2662022DKYJ003), China Postdoctoral Science Foundation (2025M783038), and the earmarked fund for CARS-41. Data from transmission electron microscopy (Thermo Talos L120C) were acquired at the public instrument center of the college of Animal Science & Technology and College of Veterinary Medicine at Huazhong agricultural university. We would be grateful to Yingying Xu for her support of sample preparation and data acquisition. We also thank the National Key Laboratory of Agricultural Microbiology Core Facility for assistance with the AKTA pure 25M system (GE Healthcare), and we are grateful to Shaoran Zhang for his help with instrument operation.

## AUTHOR CONTRIBUTIONS

P.T., M.L., and E.S. designed the experiments. M.L., E.S., S.W., H.S., S.K., G.L., Y.L., Z.X., X.Z., J.H., and Z.D. conducted the experiments. M.L., E.S., and Y.H. analyzed the data. P.T., L.S., and V.B.R. supervised the study. P.T., M.L., and E.S. wrote the manuscript. P.T. and V.B.R. revised and polished the manuscript.

