## Supplementary for "Periplasmic gatekeeping of phage DNA entry by rSAM enzyme-mediated maturation of an effector with His-X-Ser repeats"

**Supplementary Fig. 1**

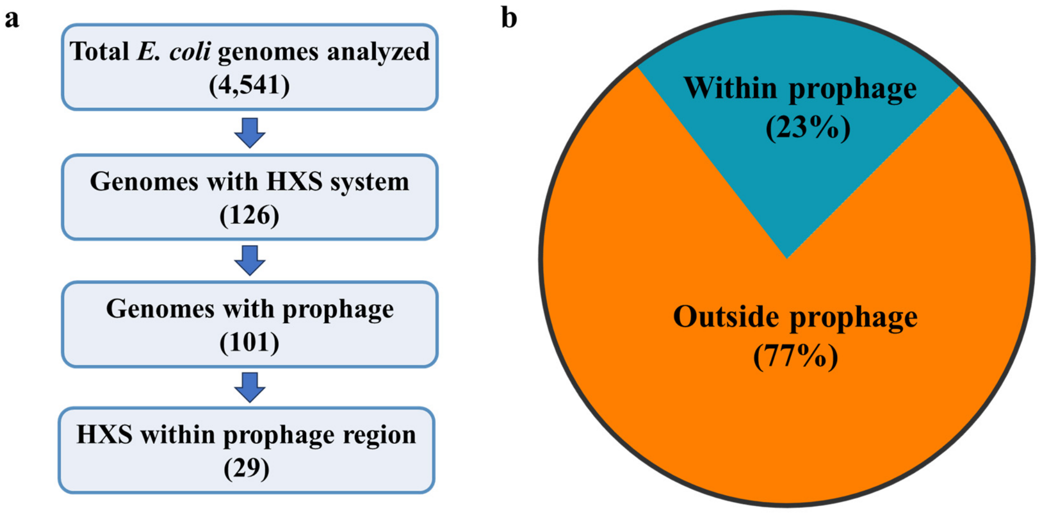


**Supplementary Fig. 1 The majority of hxsA-D operons do not overlap with prophage regions**

**a**, Schematic of the genome screening. A total of 4,541 *Escherichia coli* genomes available as of November 2025 were screened for HXS systems, identifying 126 genomes carrying HXS operons. Among these 126 genomes, 101 contained prophage regions annotated using geNomad v1.8.0, and 29 HXS systems overlapped with prophage regions. **b**, Pie chart showing the proportion of HXS systems that intersect with prophage regions (29/126, 23%) versus those that do not (97/126, 77%).

**Supplementary Fig. 2 Phage T7 escape from the HXS system**

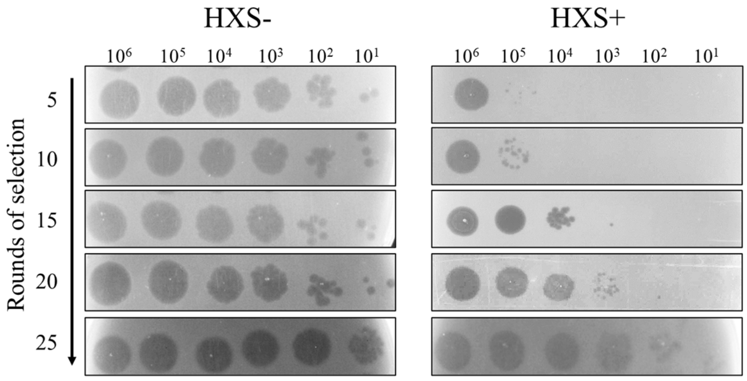


T7 was repeatedly passaged on HXS-expressing *E. coli* BL21(DE3) cells as described in Methods. Escape-evolved T7 phages were serially diluted tenfold and spotted on lawns of *E. coli* BL21(DE3) carrying an empty vector (HXS-) or the HXS system (HXS+). Plates were incubated at 37 °C overnight and imaged to assess plaque formation.

**Supplementary Fig. 3**

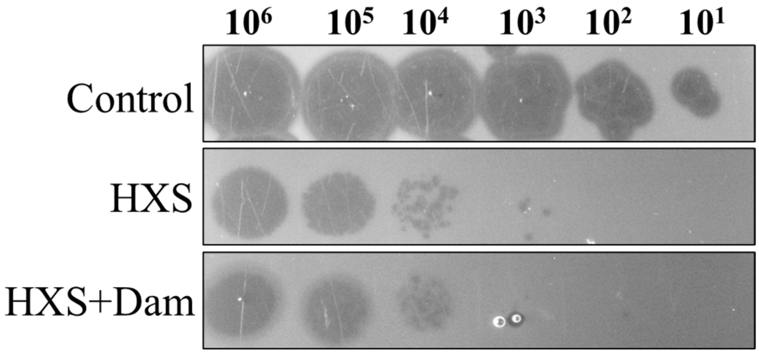


**Supplementary Fig. 3 Overexpression of Dam methylase does not affect the antiphage activity of the HXS system**

*E. coli* BL21 cells carrying HXS were transformed with a plasmid expressing *E. coli* Dam methylase or the corresponding empty vector. Ten-fold serial dilutions of T7 were spotted onto lawns of the indicated strains and incubated overnight at 37 °C. Antiphage activity was evaluated by comparing the highest dilution at which plaques were observed on HXS+ versus HXS- lawns.

**Supplementary Fig. 4**

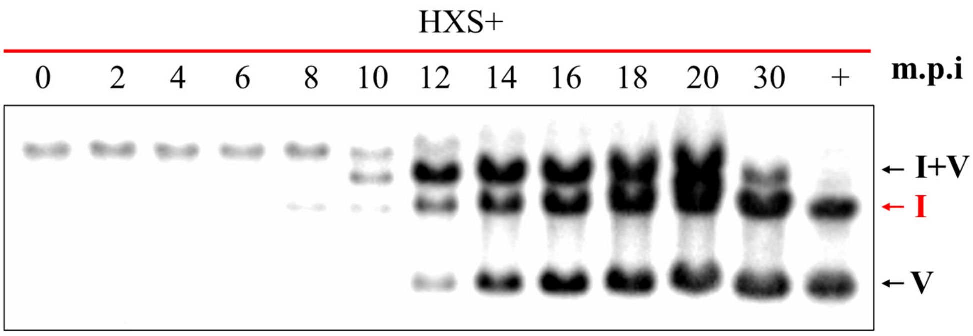


**Supplementary Fig. 4 Southern blot analysis of DNA translocation of T7 mutant in the presence of HXS system**

*E. coli* BL21 cells overexpressing Dam methylase were transformed with the plasmid expressing the HXS system (HXS+), and then infected with the HXS-escape T7 mutant. At different time points after infection, the infection mixtures were collected to isolate phage DNA, which was then digested with Dpn I. After agarose gel electrophoresis, the DNA was transferred to nylon membranes and analyzed by Southern blot using the fragment I-specific and fragment V-specific probes.

**Supplementary Fig. 5**

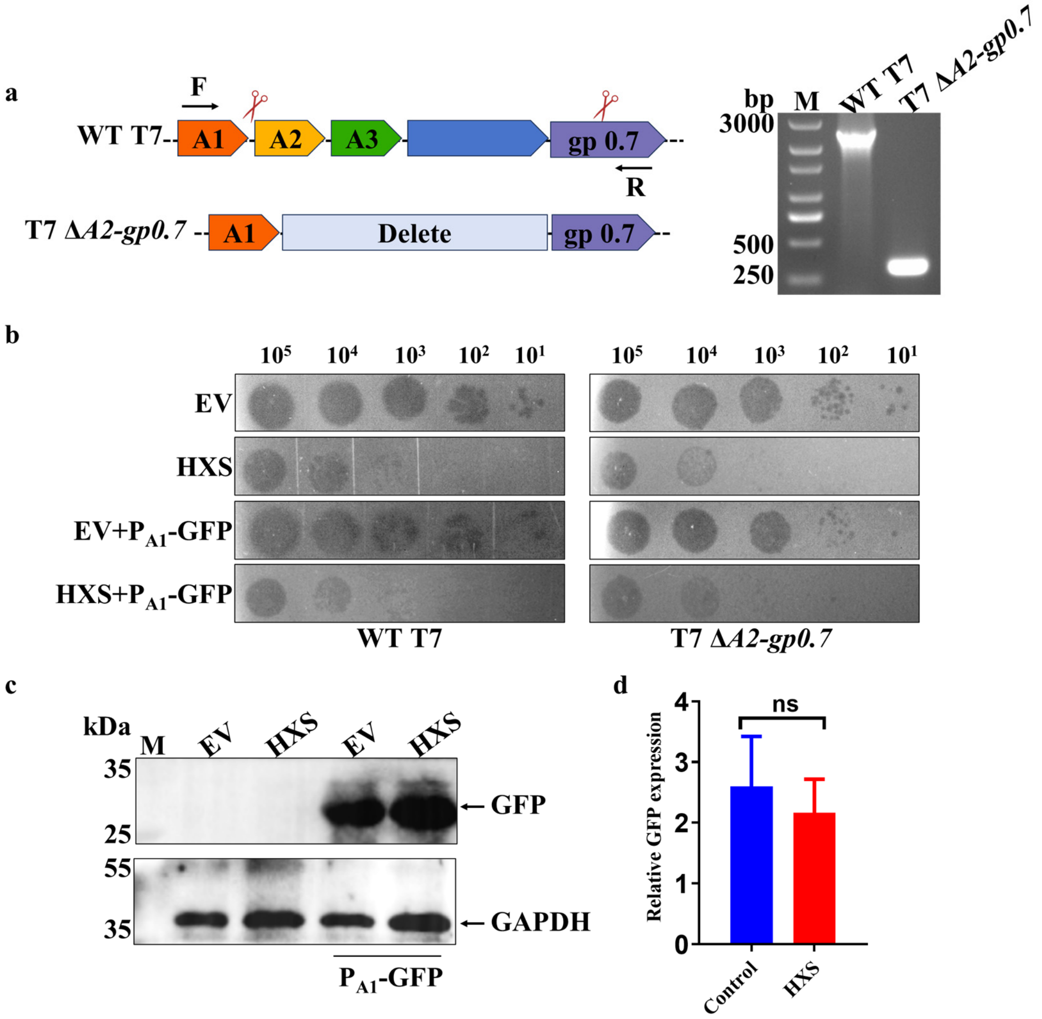


**Supplementary Fig. 5 HXS restriction is independent of $\sigma^{70}$-dependent early transcription in T7**

**a**, Schematic showing the deleted regions in the T7 *ΔA2-gp0.7* phage, with deletion sites indicated by scissors. PCR primers (F and R) used to analyze the deletion mutants are indicated. PCR products confirm that the mutants carry deletions of the expected size, with WT T7 phage used as a control. **b**, Phage challenge assay. A plasmid encoding GFP under the control of the $\sigma^{70}$ A1 promoter was introduced into *E. coli* MG1655 strains harboring or lacking the HXS system. Tenfold serial dilutions of wild-type T7 or an A2/A3-deleted T7 mutant (T7 Δ*A2-gp0.7*) were spotted onto lawns of the indicated strains and incubated overnight at 37 °C. **c**, Western blot analysis of GFP expression under the control of $\sigma^{70}$ promoter A1 in the presence of the HXS system. The empty vector (EV) was used as a control. **d**, GFP protein levels were quantified from the blots (left) by band intensity and normalized to GAPDH. Data represent the mean ± SD from three independent experiments.

### Supplementary Fig. 6

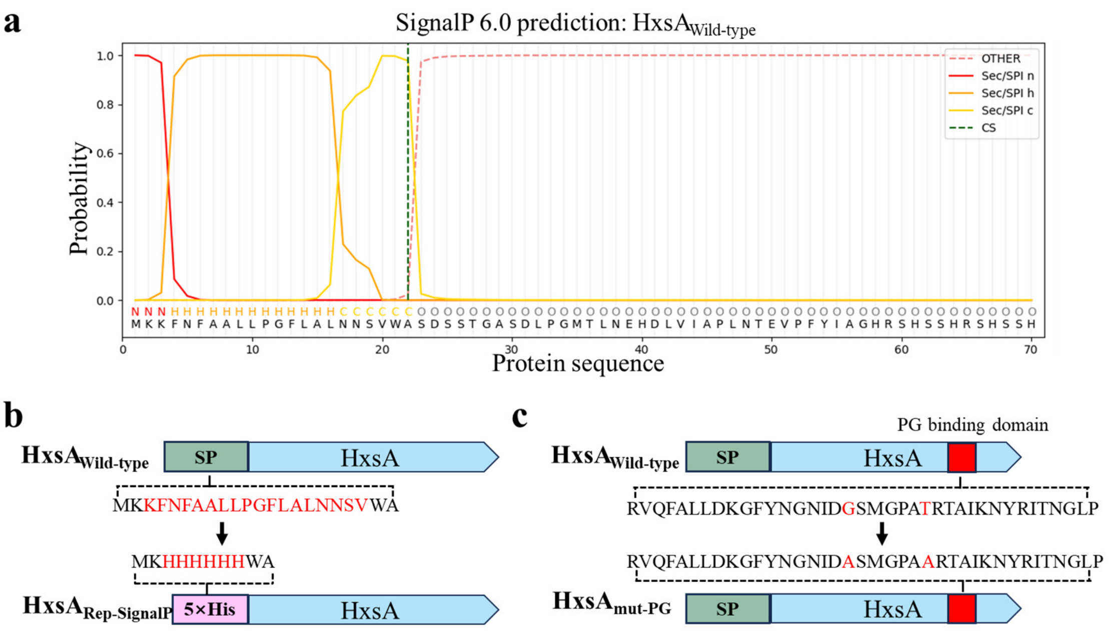


**Supplementary Fig. 6 Signal peptide prediction and engineering of HxsA variants**

**a**, The N-terminal sequence of HxsA was analyzed using SignalP, revealing a predicted Sec-type signal peptide with a putative signal peptide region and a consensus signal peptidase cleavage site (vertical line). Regions corresponding to the signal peptide and mature HxsA are indicated. **b-c**, Design of $HxsA_{Rep\text{-}SignalP}$ and $HxsA_{mut\text{-}PG}$ constructs. In $HxsA_{Rep\text{-}SignalP}$, the native HxsA signal peptide was replaced by a His tag; in $HxsA_{mut\text{-}PG}$, conserved residues within the PG-binding domain were mutated to disrupt peptidoglycan association. These constructs were used to dissect the role of signal peptide-mediated export and PG binding in HXS defense.

**Supplementary Fig. 7**

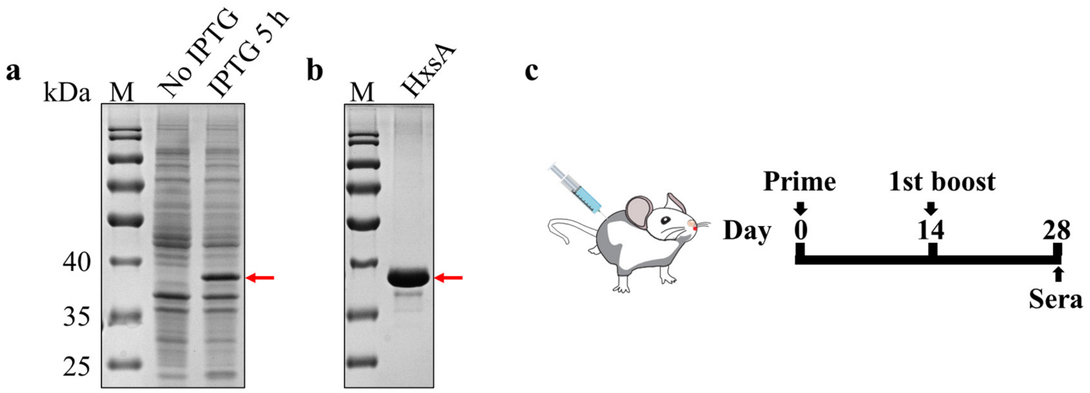


**Supplementary Fig. 7 Preparation of HxsA-specific polyclonal antibodies**

**a-b**, Expression and purification of recombinant HxsA. The hxsA gene was cloned into an inducible expression vector and expressed in *E. coli*. Cell lysates (**a**) and Ni-NTA affinity-purified fractions (**b**) were analyzed by SDS-PAGE, showing a prominent band at the expected molecular weight of full-length HxsA. **c**, Generation of anti-HxsA polyclonal serum. Mice were immunized with purified HxsA protein according to the schedule described in Methods. Sera collected after the final boost were tested by western blot against native HxsA from *E. coli* expressing HXS, confirming robust and specific anti-HxsA reactivity.

**Supplementary Fig. 8**

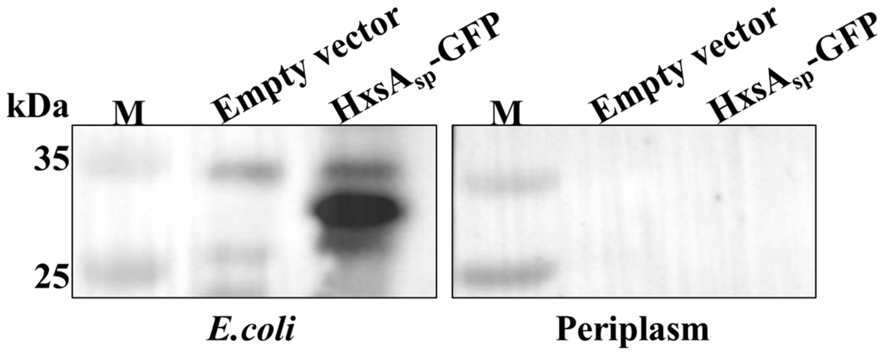


**Supplementary Fig. 8 Functional analysis of the HxsA signal peptide**

The predicted HxsA signal peptide was fused to the N-terminus of GFP (HxsA$_{SP}$-GFP). Whole-cell lysates and periplasmic fractions were prepared from *E. coli* cells expressing GFP and analyzed by western blot using anti-GFP antibodies.

**Supplementary Fig. 9**

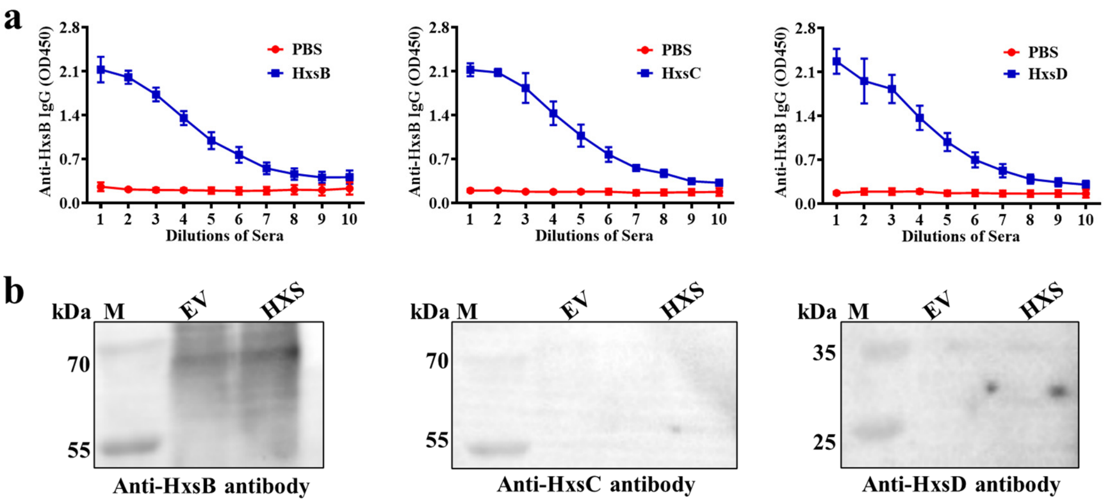


**Supplementary Fig. 9 Limited expression of HxsB, HxsC, and HxsD in cells expressing the HXS system**

**a**, Mice were immunized twice with purified His-tagged HxsB, HxsC, or HxsD at a 2-week interval, and sera collected 14 days after the second immunization were tested by ELISA on plates coated with the same proteins to determine antigen-specific antibody responses. X-axis numbers represent 2-fold serial dilutions of serum starting at 1:3,200, with number 1 corresponding to 1:3,200. **b**, Endogenous HxsB, HxsC, and HxsD proteins in whole-cell lysates were detected by immunoblotting with anti-HxsB, anti-HxsC, or anti-HxsD antibodies.

**Supplementary Fig. 10**

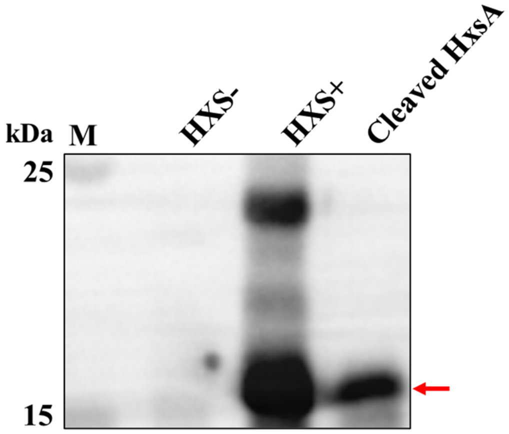


**Supplementary Fig. 10 SDS-PAGE analysis of purified cleaved HxsA protein**

*E. coli* cells expressing the HXS system were collected and analyzed by SDS-PAGE. The band corresponding to the cleaved HxsA was excised from the SDS-PAGE gel, and the protein was recovered by electrophoresis. The recovered sample was analyzed by western blot using HxsA-specific antibodies. The cleaved HxsA proteins were indicated by the red arrow. *E. coli* cells containing the HXS system (HXS+) or empty vector (HXS-) were used as controls.

**Supplementary Fig. 11**

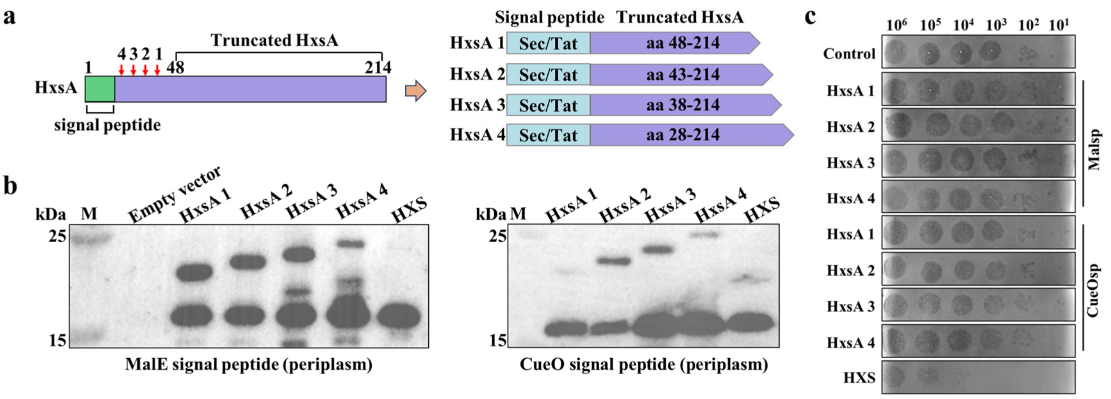


**Supplementary Fig. 11 Periplasmic targeting of HxsA truncations is insufficient for antiphage defense**

**a**, Design of N-terminally truncated HxsA variants. Four truncations of HxsA were generated starting at each predicted chymotryptic cleavage site within residues 23-47, and each was fused to either a Tat (MalE) or Sec (CueO) signal peptide, yielding eight constructs that direct truncated HxsA to the periplasm. **b**, Periplasmic localization of HxsA truncations. Periplasmic fractions from cells expressing each construct were analyzed by western blot using anti-HxsA antibodies. **c**, Evaluation of antiphage activity. Plaque assays with T7 were performed on strains expressing each periplasmic truncation construct.

**Supplementary Fig. 12**

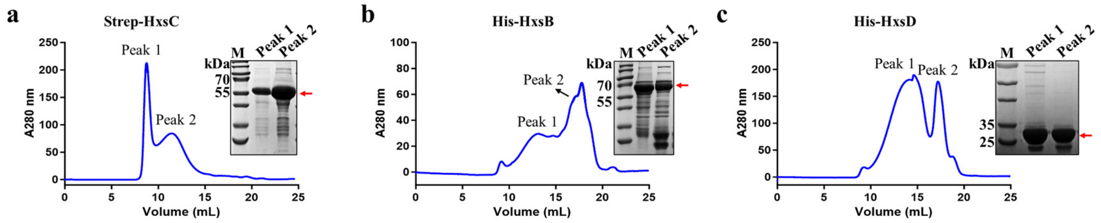


**Supplementary Fig. 12 Expression and purification of HxsB, HxsC, and HxsD proteins**

**a-c**, Recombinant HxsB and HxsD were purified by HisTrap affinity chromatography, whereas HxsC was purified using Strep-tag affinity chromatography, followed by size-exclusion chromatography. Chromatograms and SDS-PAGE analysis of peak fractions demonstrate the purity and oligomeric state of HxsB, HxsC, and HxsD used for in vitro interaction assays.

**Supplementary Fig. 13**

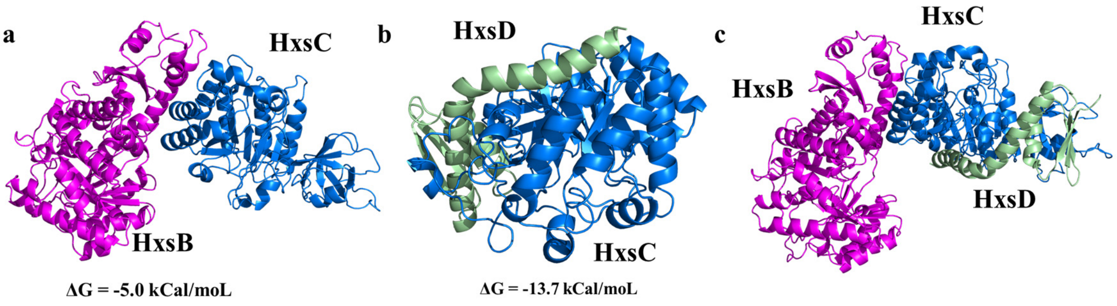


**Supplementary Fig. 13 Predicted structural interfaces between HxsB, HxsC and HxsD**

**a-c**, AlphaFold 3 model of the HxsC-HxsB (**a**), HxsC-HxsD (**b**), and HxsB-HxsC-HxsD (**c**) heterocomplex. HxsB (red), HxsC (blue), and HxsD (green) are shown. Interface solvation free energies were analyzed with PDBePISA.

**Supplementary Fig. 14**

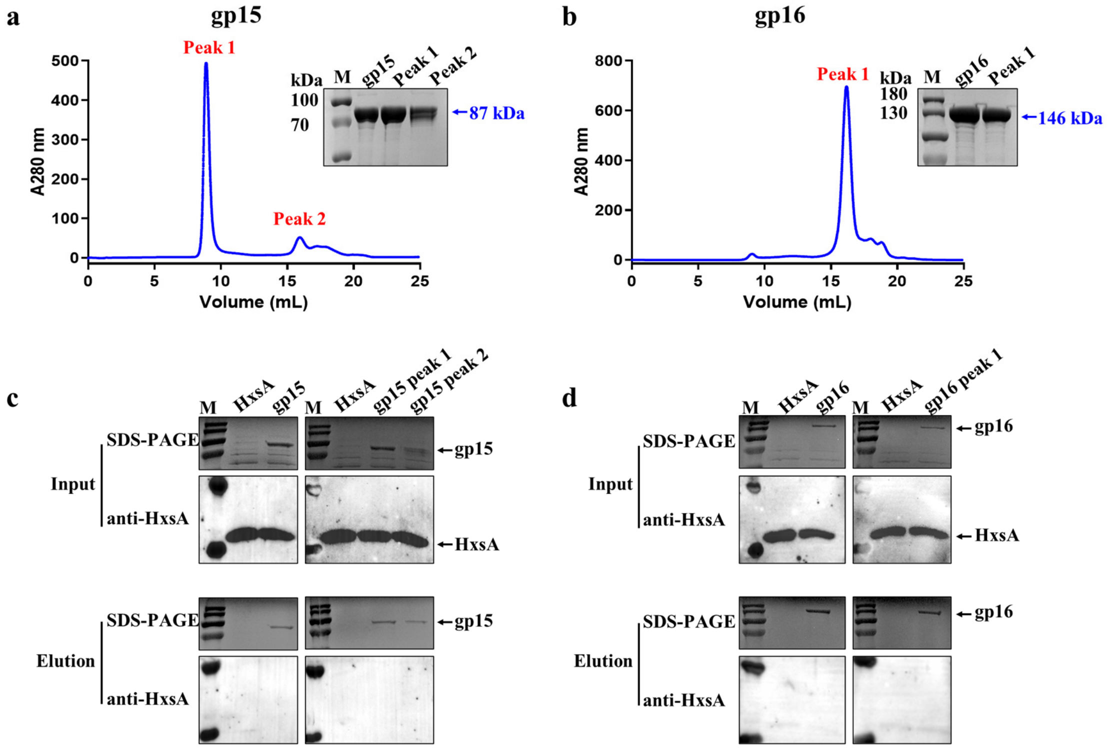


**Supplementary Fig. 14 The cleaved HxsA protein does not directly interact with gp15 or gp16**

**a-b**, Purification of gp15 and gp16. Recombinant gp15 and gp16 were purified using HisTrap affinity chromatography followed by size-exclusion chromatography, and the purified proteins were analyzed by SDS-PAGE. **c-d**, Pull-down assays testing interactions between cleaved HxsA and T7 gp15/gp16. Purified His-tagged gp15 or gp16 was incubated with periplasmic proteins containing cleaved HxsA and captured on Ni-NTA resin. Bound fractions were analyzed by SDS-PAGE and western blot using anti-HxsA serum.

**Supplementary Table 1 Taxonomic classification and GenBank accession of 59 coliphages**

| Genus | Phage ID | GenBank accession number |
|---|---|---|
| Christensenvirus | 97 | PX794797 |
| Teseptimavirus | 45 | PX794793 |
| | 57 | PX794759 |
| | 59 | PX794760 |
| | 108 | PX794799 |
| | T7 | V01146.1 |
| Tunavirus | 20 | PX794792 |
| Warwickvirus | 5 | PX794779 |
| | 6 | PX794766 |
| | 8 | PX794780 |
| | 10 | PX794757 |
| | 19 | PX794781 |
| | 21 | PX794782 |
| | 29 | PX794769 |
| | 31 | PX794805 |
| | 64 | PX794787 |
| | 66 | PX794775 |
| | 78 | PX794776 |
| | 85 | PX794790 |
| | 95 | PX794796 |
| Tequintavirus | 77 | PX794788 |
| | T5 | AY543070 |
| Hanrivervirus | 3 | PX794765 |
| | 7 | PX794802 |
| | 9 | PX794803 |
| | 23 | PX794804 |
| | 24 | PX794783 |
| | 26 | PX794767 |
| | 28 | PX794768 |
| | 30 | PX794758 |
| | 38 | PX794772 |
| | 42 | PX794786 |
| | 51 | PX794794 |
| | 62 | PX794773 |
| | 63 | PX794774 |
| | 73 | PX794761 |
| | 86 | PX794778 |
| | 90 | PX794791 |
| | 110 | PX794807 |
| | 114 | PX794800 |
| | 116 | PX794763 |
| | 117 | PX794801 |

**Supplementary Table 1 (continued)**

| | | |
|---|---|---|
| Vequintavirus | 1 | PX794764 |
| Dhillonvirus | 32 | PX794770 |
| | 33 | PX794771 |
| | 34 | PX794754 |
| | 35 | PX794755 |
| | 36 | PX794784 |
| | 41 | PX794785 |
| | 55 | PX794756 |
| | 70 | PX794795 |
| | 75 | PX794808 |
| | 76 | PX794762 |
| | 80 | PX794806 |
| | 81 | PX794777 |
| | 106 | PX794798 |
| Tequatrovirus | T4 | NC_000866 |
| | 83 | PX794789 |
| Punavirus | P1 | NC_005856.1 |

**Supplementary Table 2 Sequence alignment of the mutant and wild-type phage genomic DNAs identified seven mutation sits**

| No. | Position-start | Position-end | Mutation Type | Sequence changes | Amino acid changes | Mutant genes |
|---|---|---|---|---|---|---|
| 1 | 28,434 | 28,439 | deletion | Δcaaggt | ΔRS | gene 15 |
| 2 | 28,628 | 28,628 | point mutation | g>a | E>K | gene 15 |
| 3 | 29,451 | 29,451 | point mutation | a>c | D>A | gene 15 |
| 4 | 32,870 | 32,870 | point mutation | g>a | G>D | gene 16 |
| 5 | 33,231 | 33,231 | point mutation | c>a | F>L | gene 16 |
| 6 | 157 | 158 | insertion | +c | - | - |
| 7 | 8,957 | 8,957 | point mutation | a>c | - | gene 2 |

Note: The Δcaaggt indicated the deletion of 6 base pairs (caaggt) and the ΔRS indicated a deletion of 2 amino acids (RS).

**Supplementary Table 3 Amino acid sequence of the HXS system**

| **Proteins** | **Sequence** |
|---|---|
| HxsD | MAMWEKILQKNLYSEWVIRNTLYWMTPVSPWKLEENTSSWIIFFEIEDPECEFEFERLL NDFSLREKLHHQTGQLRDAIISKVLRSIDDRLAK |
| HxsB | MRLMPFNFDRMPDGRVFISNLAGFHHFVGEQEFFDLVNGHISLEQSSSLESKLFVCGEES SSVTPHALSSAFAKRLMNELAVRPIFMIVPTLRCDHTCKYCQVSRASVTAYGYDLNPELI PQIVNTIKKLGTPPYKIEVQGGEPLLRFDLVQSIYHECEISLGSGAFEMVIATSLSLLDESI LSWVKERNITFSVSLDGNEAVHNKNRLLTDFQAHNKAVTGIRKITEELGANRVATVTTV TKELTKEPASIVDAHLSLGLTDMFIRPVSPYGFAQKQSFTFSMPEYFAFYKELMQEVLIQ NEKGIPVIEHSAAIHLKRIFNPGFSGYADLKSPSGVVLNCILFNYDGKVYGSDESRMLQK VNPEADFSAGEFASLSFISNEFYRSALSSSFNFAMPGCDTCAYQPFCGADPCQNISVHGE PVGDKSRSTFCQYHKGMFRFLLNEISQDGPMAKMLKGWVYV |
| HxsC | MSEVLRNDIFRFTADLPVQQGFYRLCKSKPENPQFYLPNLLVSETELDSRLPAYFADYVV STNLFNSIEDGDIGIVNNGNMIRVILSRRANHNTVLVTERCNNSCLFCSQPPKTGNDDWL LNQSALAIASFALNGVVGVSGGEPLLYGEDFLQFLDFIIENSPETALHVLTNGRKFADVSF TEQMKERSEKLKITFGIPLYSSRSSVHDYLVGSEGAFDETVRGLINAGNSGINIELRIIPTL ANYMELDKIIEFAGRVFSNINQISLMGLESIGWARKNWSSIFIEHDSYSEKILSVIGTAQRS GIPLTIFNYPLCHLPERAWRFATQSISDWKNYYPKECDECTQKSSCAGYFSSSKGRFHQP PRPIL |
| HxsA | MKKFNFAALLPGFLALNNSVWASDSSTGASDLPGMTLNEHDLVIAPLNTEVPFYIAGHR SHSSHRSHSSHRSSSGGGYYGGSTPYYPKTYSSPSSSGSSSSGTSSSSSSSSIRSLRSNDMD TSTTTNSNGTNGSNGNNRSSSGVASDTEKRKRLIMRVQFALLDKGFYNGNIDGSMGPAT RTAIKNYRITNGLPTPARETLDTQLLNSLNILAR |

**Supplementary Table 4 Sequences of the spacers used for T7 phage genome editing**

| No. | Target edit sites | Spacer sequences |
|---|---|---|
| 1 | I | actcatagctcattacctccccg |
| 2 | II | atcttctgcataacgtcatcgtc |
| 3 | III | ttcattgacttcatggaatcgtc |
| 4 | IV | attcggacctatccatcactatg |
| 5 | V | ccacacagtgtcctgattgcgtc |

**Supplementary Table 5 Primers used to amplify T7 phage DNA**

| Primers | Sequences (5′-3′) | Length |
|---|---|---|
| T7 - southern - I - F | tgaagtgtctcacggctgac | 899 bp |
| T7 - southern - I - R | tcagcccattaacattgcgtc | |
| T7 - southern - V - F | gttacgcgctgagtacgatgag | 867 bp |
| T7 - southern - V - R | aactgcatggactcacggag | |
